# A Fibronectin (FN)-Silk 3D Cell Culture Model as a Screening Tool for Repurposed Antifibrotic Drug Candidates for Endometriosis

**DOI:** 10.1101/2024.10.05.616776

**Authors:** Sarah Teworte, Mark C. Behrens, Mona Widhe, Lukas-Adrian Gurzeler, My Hedhammar, Paola Luciani

**Affiliations:** Pharmaceutical Technology Research Group, Department of Chemistry, Biochemistry and Pharmaceutical Sciences, University of Bern, Freiestrasse 3, CH-3012 Bern, Switzerland; RNA Biology Research Group, Department of Chemistry, Biochemistry and Pharmaceutical Sciences, University of Bern, Freiestrasse 3, CH-3012 Bern, Switzerland; Division of Protein Technology, School of Biotechnology, KTH Royal Institute of Technology, AlbaNova University Center, Stockholm SE-106 91, Sweden

**Keywords:** 3D cell culture, endometriosis, fibrosis, drug repurposing, FN-silk, pirfenidone

## Abstract

This study advances sustainable pharmaceutical research for endometriosis by aligning with the UN Sustainable Development Goals on health, gender equality, and responsible consumption in developing *in vitro* 3D cell culture models of endometriotic pathophysiology. Fibrosis is a key aspect of endometriosis, yet current models to study it remain limited, especially in 3D. This work aims to bridge the translational gap between *in vitro* fibrosis research and preclinical testing of non-hormonal drug candidates. When grown in a 3D matrix of sustainably produced silk protein functionalized with a fibronectin-derived cell adhesion motif (FN-silk), endometrial stromal and epithelial cells respond to transforming growth factor beta-1 (TGF-β1) in a physiological manner as probed at the mRNA level. For stromal cells, this response to TGF-β1 is not observed in spheroids, while epithelial cell spheroids behave similarly to epithelial cell FN-silk networks. Pirfenidone, an antifibrotic drug approved for the treatment of idiopathic pulmonary fibrosis, reverses TGF-β1-induced upregulation of mRNA transcripts involved in fibroblast-to-myofibroblast transdifferentiation of endometrial stromal cells in FN-silk networks, supporting the drug’s potential as a repurposed non-hormonal therapy for endometriosis. This study demonstrates how a sustainable approach – from project conceptualization to material selection – can be integrated into pharmaceutical research for women’s health.

**Table of contents:** This paper presents *in vitro* 3D cell culture models of fibrosis in endometriosis. Endometrial stromal and epithelial cells cultured in networks of silk protein functionalized with a fibronectin-derived cell adhesion motif showed physiological-like fibrotic behavior. Pirfenidone was able to reverse fibrosis of endometrial stromal cells *in vitro*, demonstrating this model’s suitability as a screening tool for antifibrotic drugs for endometriosis.

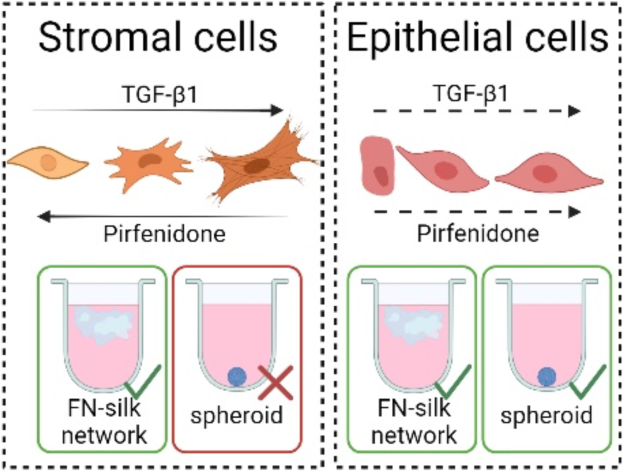

## 1. Introduction

Gender equality is a fundamental right,[1] and true empowerment of women can only be achieved when advancements in women’s health are prioritized, adequately funded, and made visible.[2–6] Despite affecting 10% of women, treatments for endometriosis remain underfunded and under-researched.[4,7,8] This work addresses the urgent need for pharmacological therapies for endometriosis, aligning with Sustainable Development Goal (SDG) #3 (Good Health and Well-Being) and SDG #5 (Gender Equality),[9] illustrating how these SDGs can be integrated into preclinical research.[2] We emphasize the importance of sustainable research practices, including the use of non-animal-derived materials, and hereby present novel 3D *in vitro* models of endometriotic stroma and epithelium that model fibrosis using a recombinant silk protein functionalized with the arginine-glycine-aspartate (RGD) cell adhesion motif from fibronectin (FN-silk). By adopting this sustainable material, we embrace our responsibility as scientists to address SDG #13 (Responsible Consumption and Production),[9] particularly regarding the materials used in our laboratories. FN-silk networks enable *in vitro* modelling of fibrosis in endometriosis and testing of an antifibrotic repurposed drug candidate for endometriosis, aiming to reduce reliance on animal experiments and improve the clinical translatability of our research.

There is ample histopathological evidence for dense, fibrous tissue in and around endometriotic lesions,[8,10–17] yet the role of fibrosis in the pathophysiology of endometriosis is only just beginning to be emphasized by the research community,[10–14] with Viganò *et al.* even suggesting the definition of endometriosis be changed to “a fibrotic condition in which endometrial stroma and epithelium can be identified”.[11] During repeated injury caused by the shedding of ectopic endometrium in response to a withdrawal of progesterone – a bleeding internal wound[18] – the body’s healing response becomes pathological, causing extracellular matrix (ECM) proteins to be deposited recurrently in an attempt to heal tissue damaged by the decidualizing endometriotic lesion.[13,19–22] This process of repeated tissue injury and repair leads to scarring, fibrosis, and, in severe cases, adhesions.[11–13,23] Encouragingly, recent preclinical studies in non-human primate and murine models have used the presence and extent of fibrotic tissue as an indicator of disease progression against which to evaluate the efficacy of drug candidates.[24,25] When developing anti-fibrotic pharmacological treatments for endometriosis, the biggest challenges are how to reverse or prevent fibrosis, and how to reach fibrotic endometriotic lesions in the pelvis and on the peritoneum.[11,26]

Cellular mechanisms involved in the fibrogenesis of endometriosis are fibroblast-to-myofibroblast transdifferentiation and epithelial-to-mesenchymal transition[14,18,19,27–31] (**Figure 1A**). On the one hand, fibrosis in endometriosis is driven by stromal fibroblasts, which transdifferentiate into myofibroblasts upon exposure to cytokines such as TGF-β1,[12,16,32] neuropeptides,[33–35] or mechanical cues.[36–39] In fibrosis, secreted ECM proteins accumulate over time, forming fibrotic lesions, which disrupt the tissue architecture and are at least in part responsible for typical symptoms of endometriosis such as infertility, chronic pelvic pain, and dyspareunia.[8,11,13,15] On the other hand, epithelial-to-mesenchymal transition enhances cell migration and adhesion to the peritoneum, thereby increasing the invasive capacity of endometriotic epithelial cells.[30,40–43] Unlike other fibrotic diseases, fibrosis in endometriosis correlates with higher pain scores.[10,33,44]

**Figure 1.**
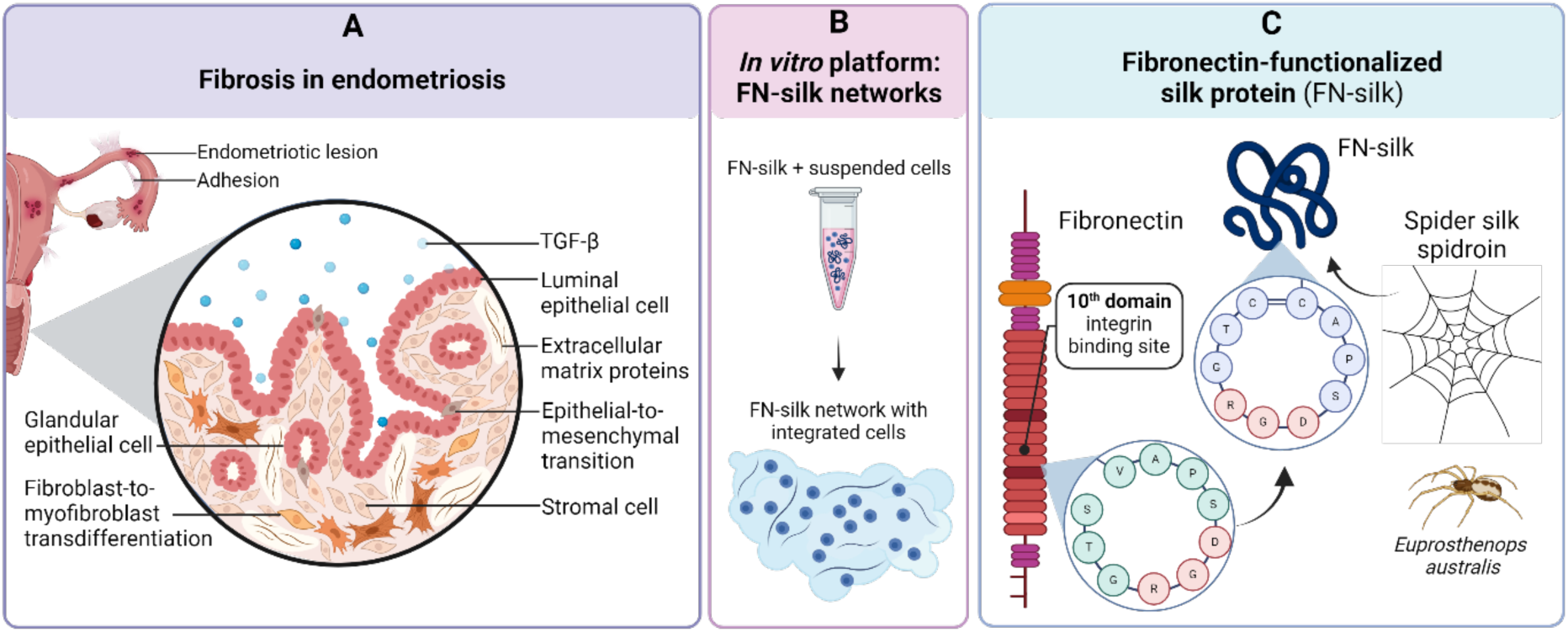
**A** Schematic illustration of TGF-β-modulated fibrotic cellular processes in an endometriotic lesion, including fibroblast-to-myofibroblast transdifferentiation, deposition of extracellular matrix proteins, and epithelial-to-mesenchymal transition (adapted from[10,12,45]). **B** Schematic illustration of the FN-silk network platform with integrated cells pioneered by Johansson, Widhe *et al.*[46] and Collodet *et al.*[47]. **C** Schematic illustration of FN-silk, a recombinant spider silk protein genetically functionalized to include the arginine-glycine-aspartate (RGD) cell adhesion motif from fibronectin (adapted from[48–50]).

More recently, 3D models in which cells grow in a matrix have raised interest to evaluate fibrotic mechanisms, as cells are able to interact with the matrix and other cells from all sides, deposit ECM proteins, and create their own biomechanical microenvironment, mimicking endometriotic foci found in humans more closely.[10,27,38,42,45,51–56] To model endometriosis and associated fibrosis, we used a recombinantly produced spider silk protein derived from the four poly-Ala/Gly-rich repeats and the non-repetitive C-terminal domain of Major Ampullate Spidroin 1, 4RepCT, genetically functionalized to include the RGD cell adhesion motif from fibronectin (FN), known as FN-silk (Figure 1C).[46–49,57–60] FN-silk possesses the unique capability to self-assemble into biodegradable and biocompatible microfibers in aqueous, physiological-like buffers at room temperature.[46,50] FN-silk has the additional advantage of having a fully defined composition and being produced using biotechnological methods, reducing environmental impact compared to animal-origin materials. The addition of a cell adhesion motif from ECM proteins facilitates cellular attachment to and proliferation on the FN-silk protein compared to non-functionalized silk proteins.[48] Harnessing the ability of the FN-silk protein to self-assemble into fibers of β-sheet structures formed at the air-liquid interface,[50] Johansson, Widhe *et al.*[46] established a method to produce networks from FN-silk with integrated cells (Figure 1B).

Here, we explore 3D cell culture to model fibrotic pathophysiology in endometriotic stroma and epithelium, comparing FN-silk network and spheroid formats. 3D cell culture in a matrix may have advantages in modelling the fibrotic condition of endometriosis, as cells are exposed to biomechanical cues arising from cell-cell and cell-matrix contacts mimicking the ECM. We demonstrate the use of the *in vitro* models developed herein to test pirfenidone, an antifibrotic drug indicated for pulmonary fibrosis that has the potential to be repurposed for endometriosis.[61,62]

## 2. Results and discussion

### 2.1. Formation and characterization of 3D systems: FN-silk networks and spheroids

The immortalized human endometrial stromal cell (T-HESC) and epithelial 12Z cell lines were chosen because they are non-cancerous, stem from patients with leiomyoma and endometriosis, respectively, are readily available, and have been used previously, allowing us to compare results to previously published work. Both the T-HESC (endometrial stromal) and the 12Z (endometriotic epithelial) cell lines were characterized in terms of their metabolic activity and morphology when grown in monolayers, to give us hints about design parameters and a context in which to evaluate results from 3D systems (Supporting Information, **Figure S.1 and S.2**). FN-silk networks with integrated cells were produced using a method adapted from Collodet *et al.*[47] as illustrated in Figure 2A, where the FN-silk protein assembled into β-sheets during foaming. The foam transformed into a network as the sheets around the bubbles burst during the first two days after production, causing the FN-silk networks to become submerged in the culture medium. Spheroids were formed by spontaneous aggregation of cells in suspension in an ultralow attachment plate (Figure 2B). Stromal T-HESCs were seeded at a density of 10,000 cells per unit and epithelial 12Z cells were seeded at a density of 20,000 cells per unit (where one unit was an FN-silk network or a spheroid). We observed that 96 ± 7% of stromal T-HESCs and 96 ± 4% of epithelial 12Z cells seeded were integrated into the FN-silk network (mean ± SD) (Figure 2C), as measured by counting the number of cells left in the supernatant after transferring the FN-silk network to a well with fresh medium (Figure 2A).

**Figure 2.**
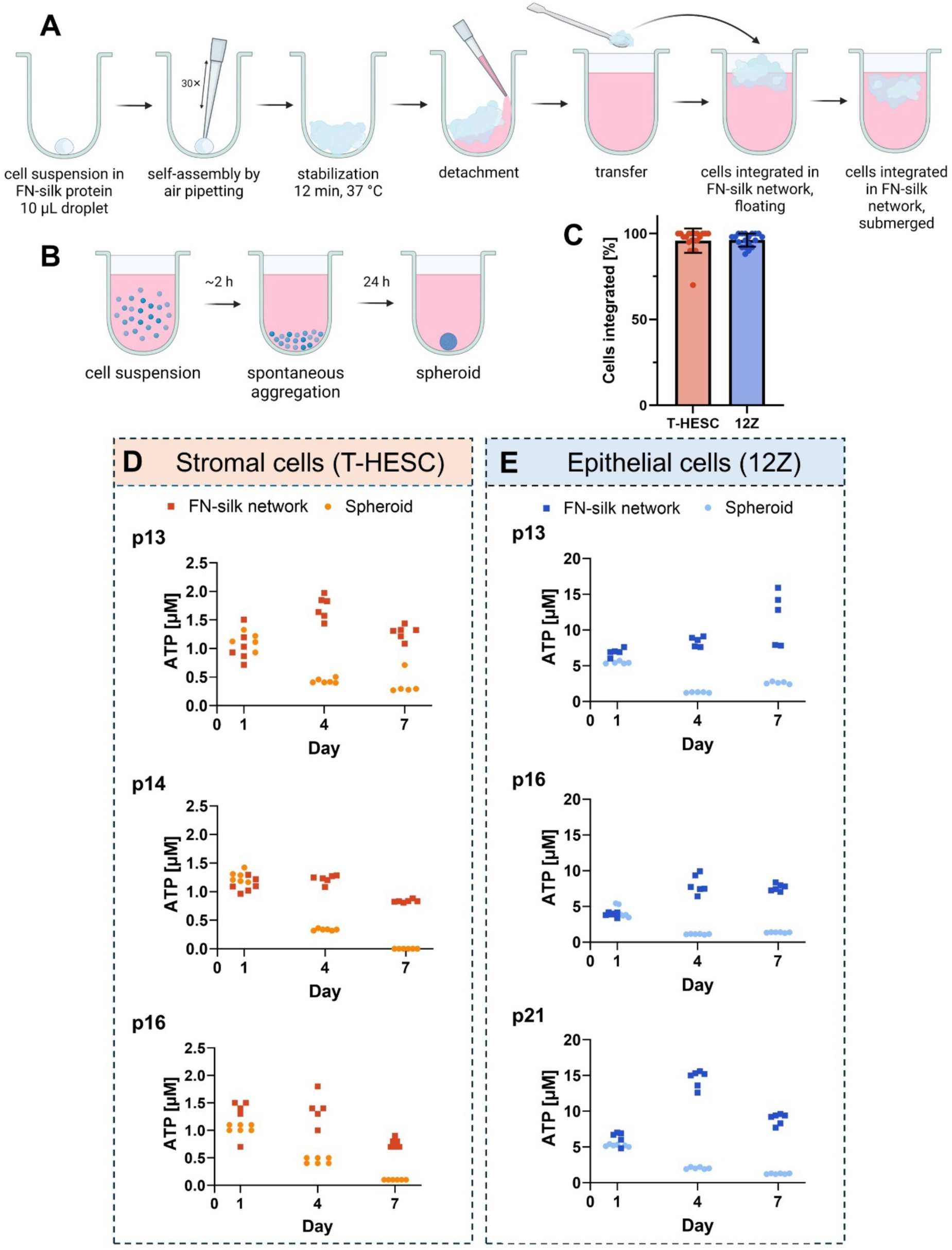
**A** Scheme depicting the workflow of the production of FN-silk networks containing cells, adapted from Collodet *et al.*[47] **B** Scheme showing the procedure for the production of spheroids. **C** Seeding efficiency of stromal T-HESC and epithelial 12Z cells in FN-silk networks as a percentage of cells seeded that were integrated into the network (individual data points and mean ± SD from N=2 independent experiments, each with n=9 or n=10 technical replicates).

To evaluate cell metabolic activity in FN-silk networks and spheroids, cells from three different passage numbers were seeded in each 3D format and the metabolic activity was measured using a CellTiter Glo^®^ 3D Viability Assay at 1, 4 and 7 days post-seeding (day 0 was the day on which systems were seeded). On day 1 post-seeding, both spheroids and FN-silk networks showed similar metabolic activity, confirming equal cell seeding (Figure 2D**, 2E**). By day 4, stromal T-HESC metabolic activity increased slightly or remained constant in FN-silk networks but dropped in spheroids (Figure 2D). By day 7, metabolic activity in FN-silk networks decreased slightly for stromal T-HESCs, while spheroids showed very low activity, suggesting cell death or dormancy (Figure 2D). For epithelial 12Z cells, we observed increased metabolic activity in FN-silk networks by day 7, with variability based on passage numbers, while spheroids showed a decline (Figure 2E). For both cell types, FN-silk networks maintained cellular metabolic activity and supported 3D proliferation, whereas spheroids exhibited a decline in metabolic activity.

The results presented here are in agreement with previously published work by Stejskalová *et al.*, who found that at day 4, endometrial stromal cell spheroids (St-T1b cell line) had lower metabolic activity than endometriotic epithelial cell spheroids (12Z cell line) produced using the hanging drop method, with both cell types seeded at 20,000 cells per spheroid.[56] These spheroids showed sprouting and migratory behavior, indicating that at least the cells at the outer shell of the spheroid remained alive and responsive to molecular cues.[56] Wendel *et al.*[63] produced 12Z spheroids seeded at 18,000 cells per spheroid using the same method described here (Figure 2B).

Metabolic activity measured using the luciferin/luciferase-based CellTiter Glo^®^ 3D Cell Viability Assay, of **D** stromal T-HESC and **E** epithelial 12Z cells cultured in FN-silk and spheroid formats seeded with cells from three different passage numbers (p), *i.e.*, N=3 independent experiments. Stromal T-HESCs were seeded at 10,000 cells per unit, and epithelial 12Z cells were seeded at 20,000 cells per unit. Individual data points with n=5 or n=6 technical replicates per format and time point.

At day 5 of Wendel *et al.*’s epithelial 12Z spheroid culture, a section cut through the center of the spheroid showed cellular expression of Antigen Kiel 67 (Ki67), a cell division marker, and no expression of cleaved Caspase 3 (cCAS3), an apoptotic marker, even at the spheroid core,[63] suggesting that 12Z cells proliferated in spheroids.[63] This contradicts the observations described here (Figure 2E), which may be due to differences in passage number. The CellTiter Glo 3D Viability Assay was compared to a manual count of cells, and the ATP levels measured were in good agreement with the number of cells counted (Supporting Information, **Figure S.4**).

To assess the distribution of cells throughout the FN-silk network and morphological changes during the culture period, FN-silk networks with green fluorescent silk were prepared. On days 1, 4 and 7, networks were fixed and the actin filaments of the cell cytoskeleton were stained with phalloidin 594, and nuclei with DAPI. Figure 3A shows that stromal T-HESC FN-silk networks contracted between culture days 1 and 4, and subsequently maintained their architecture over a 7-day culture period. Cells were uniformly distributed throughout the FN-silk matrix, with slightly more cells at the outer edges of the network (Figure 3A). The projected area of stromal T-HESC FN-silk networks was calculated from bright field images and was observed to decrease from 5.9 ± 1.2 ×10^6^ µm^2^ at day 1 to 2.3 ± 0.3 ×10^6^ µm^2^ at day 4 and 1.5 ± 0.3 ×10^6^ µm^2^ at day 7 (mean ± SD from n = 6 FN-silk networks, bright field images not shown). This reduction in projected area is attributable to the bursting of bubbles during the formation of the FN-silk network at culture days 1 and 2, after which all bubbles had burst. Subsequently, stromal T-HESCs remodeled the FN-silk network, as cells were oriented along the FN-silk fibers and contracted the FN-silk matrix. The thickness of the networks was in the range of 100−150 µm, estimated from z-stack images taken using confocal microscopy, which was thin enough to allow sufficient oxygenation of cells throughout the network.[64] Stromal T-HESCs tended to stretch along the silk fibers, similar to results reported previously on the SK-BR-3 and MDA-MB-231 breast cancer cell lines.[47]

**Figure 3.**
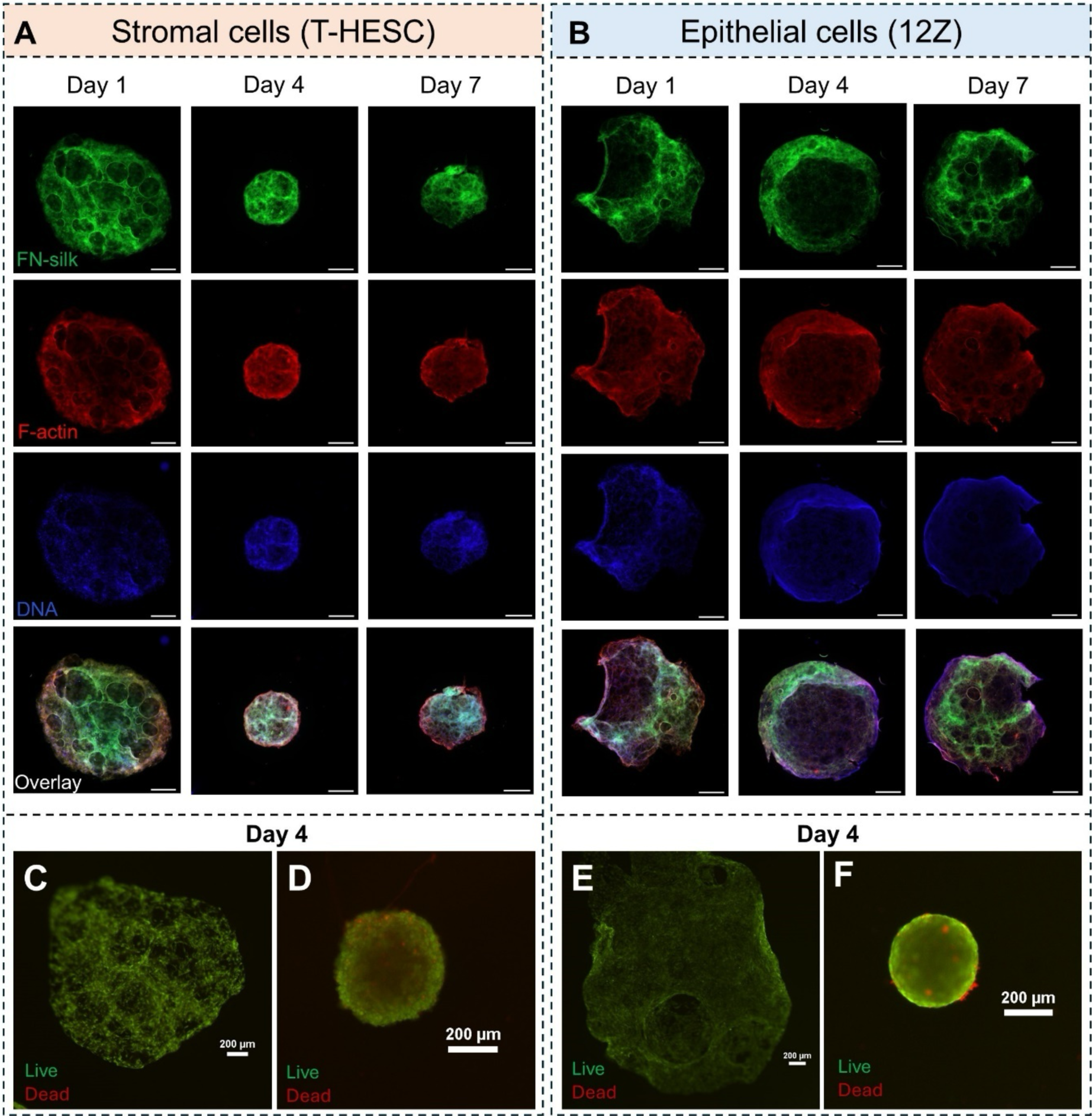
Morphology of **A** stromal T-HESC and **B** epithelial 12Z in FN-silk networks at culture days 1, 4 and 7. Single channel images of FN-silk (green), F-actin (red), DNA (blue), and overlay. Viability of **C** T-HESC in FN-silk networks, or **D** spheroids, and **E** 12Z in FN-silk networks, or **F** spheroids at culture day 4, visualized using calcein AM (green) to stain the cytoplasm of live cells and SYTOX Deep Red (red) to stain the nuclei of dead cells. Scale bars indicate 500 µm (**A, B**) or 200 µm (**C–F**).

The establishment of strong cell-matrix contacts by stromal T-HESCs was further evidenced by experiments in which cells were detached from FN-silk networks (Supporting Information, **Figure S.3**). It took relatively harsh conditions of 0.5% Trypsin-EDTA incubation for 30 min to detach cells that had been cultured in FN-silk networks for 7 days; shorter incubation periods, lower Trypsin concentrations and more gentle cell dissociation reagents were unable to detach cells from the FN-silk matrix. This indicates that cells established strong contacts with the FN-silk, mimicking interactions with the ECM. Similar to stromal T-HESCs, epithelial 12Z FN-silk networks maintained their architecture over a 7-day culture period (Figure 3B). On day 1, epithelial 12Z cells were uniformly distributed on the fibers of the FN-silk network. During proliferation in the FN-silk network, epithelial 12Z cells appear to have grown at the outer edges of the silk networks, after filling the center.

To cross-check results from the metabolic activity assay, which works by ATP measurement, we evaluated cell viability in 3D culture formats visually using a live/dead staining kit, shown in Figure 3C-F. A homogeneous distribution of live cells was observed throughout the FN-silk network, with few dead cells visible, and some empty regions corresponding to the locations of the bubbles produced during the manufacturing procedure (Figure 3C**, 3E**). Spheroids of stromal T-HESCs (Figure 3D) and epithelial 12Z cells (Figure 3F) showed mostly live and some dead cells, more than can be seen in the FN-silk network with each respective cell type. While qualitative, these images support the fact that cells in FN-silk networks had higher levels of metabolic activity, while spheroids had lower levels of metabolic activity, and suggests that this may be in part attributable to more dead cells in the spheroids. Overall, the FN-silk networks enabled cells to live and proliferate in 3D, while spheroids showed a reduction in metabolic activity and more dead cells. The FN-silk networks enabled sufficient oxygenation and more cell-cell interactions with less oxidative stress, cell death, and senescence compared to culture in a spheroid format.[65]

To characterize the architecture of the cells in FN-silk networks more closely, samples were fixed on day 4 followed by immunofluorescent staining. Figure 4A shows the morphology of stromal cells in FN-silk networks, stained for vimentin, a mesenchymal cell marker. Figure 4B shows 12Z cells stained for epithelial cell adhesion molecule (EpCAM) in an FN-silk network. Stromal and epithelial cells were uniformly distributed throughout the FN-silk matrix. Close-up images (**Figure S.6)** of the FN-silk networks show that cells were closely localized to the matrix, facilitated by strong cell-matrix interactions mediated by integrins *via* the RGD cell adhesion motif on the FN-silk.

**Figure 4.**
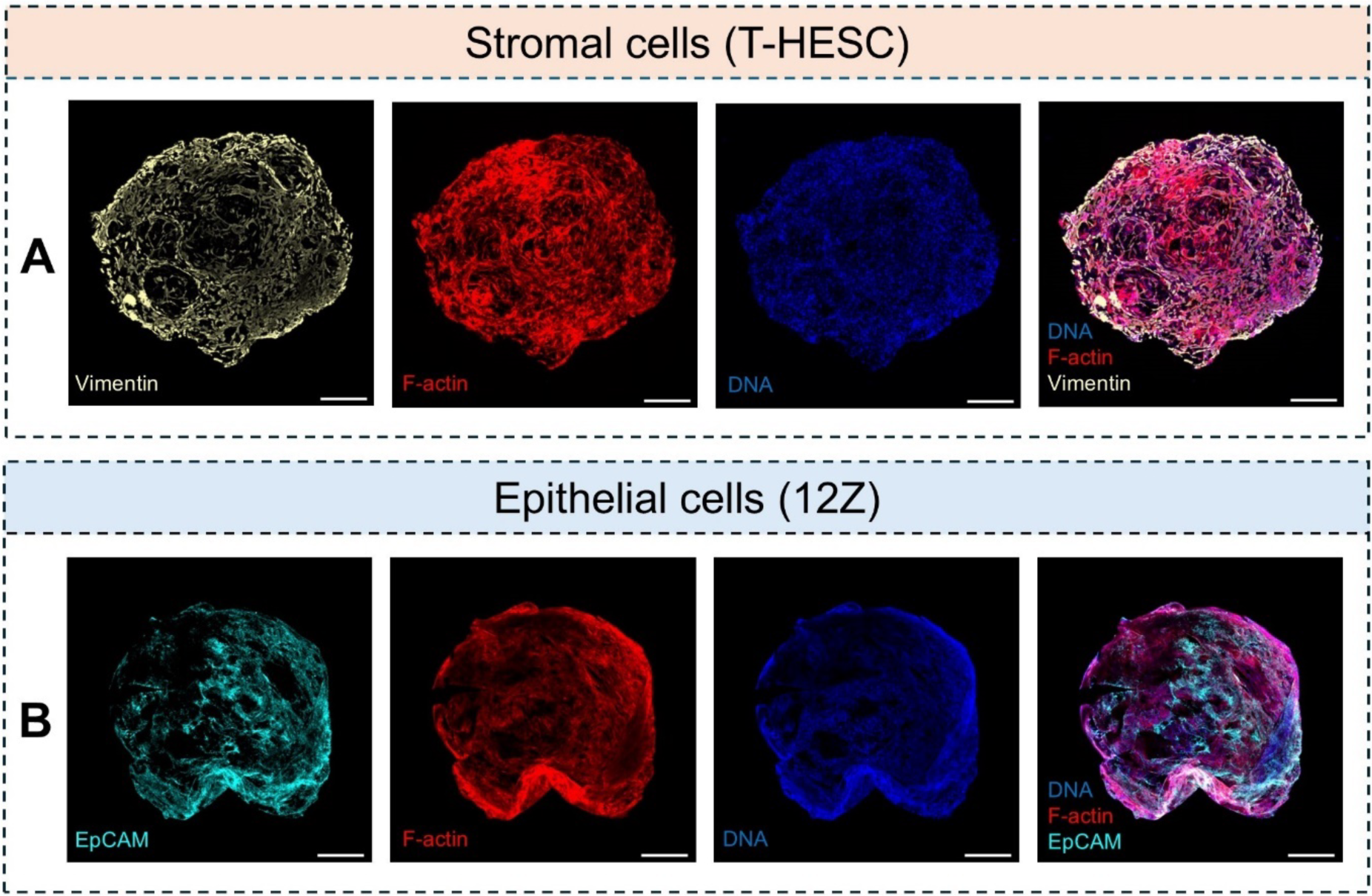
Morphology of fixed and immunostained FN-silk networks at culture day 4. **A** Stromal T-HESC FN-silk networks stained for vimentin and **B** Epithelial 12Z cell FN-silk networks stained for epithelial cell adhesion molecule (EpCAM). Scale bars indicate 500 µm.

### 2.2. Biomechanical characterization of FN-silk networks

The biomechanical properties of stromal T-HESC FN-silk networks were characterized using bioindentation, a technique that allowed us to probe the mechanical properties at the whole-network scale (in the millimeter range, Figure 5A). Figure 5B shows a representative force-depth plot of an FN-silk network at culture day 6. The force-depth curve of the FN-silk network showed a linear increase in the applied compressive force during the loading phase, with a penetration depth of 46 μm at the maximum compressive load (Figure 5B), corresponding to about 30-50% of the height of the FN-silk network. The unloading segment shows the FN-silk network’s response to the release of the compressive force, providing insights into its elastic recovery behaviour. The FN-silk network showed a relatively low gradient of the unloading curve, indicating a very elastic, low-stiffness material. The control curve (Figure 5B) shows a lower penetration depth of 5.5 μm and a steeper unloading curve gradient, indicating a higher-stiffness material, in this case, glass. The compressive elastic modulus of stromal T-HESC FN-silk networks was in the range of 0.41–0.57 kPa at culture day 3 and 0.40–0.52 kPa at culture day 6 (Figure 5C), indicating that no measurable differences in the compressive elastic modulus were detected at different culture periods. This suggests that the primary elastic properties stem from the FN-silk matrix itself, rather than any action of cells in the network. Figure 5D shows the estimated elastic modulus *EIT* of stromal T-HESC FN-silk networks. We estimated the elastic modulus of stromal T-HESC FN-silk networks to lie in the range of 1.8–4.6 kPa (Figure 5D). This elastic modulus is similar to that reported for endometrial tissue and materials used to model healthy/low stiffness conditions of the endometrium and endometriosis *in vitro*,[16,38,42,52,66–69] underscoring the ability of FN-silk to provide a physiologically relevant matrix in terms of the biomechanical environment that cells reside in.

**Figure 5.**
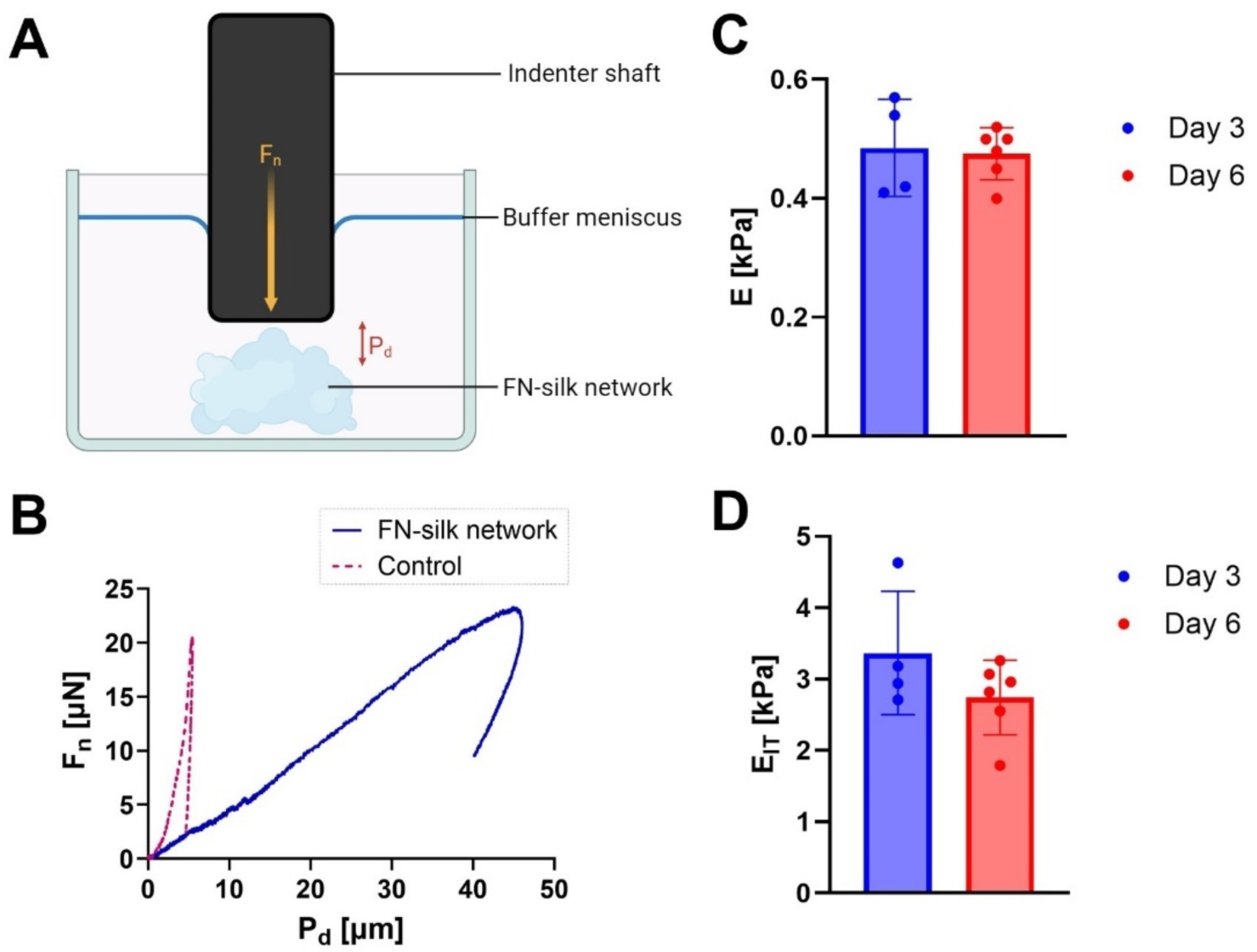
Biomechanical characterization of FN-silk networks. **A** Schematic of the bioindentation method, showing the indenter shaft applying a compressive force Fn to the FN-silk network, causing the FN-silk network to be compressed by a penetration depth Pd. **B** Representative force-depth plot of a stromal T-HESC FN-silk network and control (glass bottom well containing PBS). **C** Compressive elastic modulus *E* and **D** elastic modulus *EIT* of stromal T-HESC FN-silk networks at culture days 3 and 6, estimated using the Oliver & Pharr method.[70] Individual data points and mean ± SD of n = 2 FN-silk networks at day 3, and n = 3 FN-silk networks at day 6. Each FN-silk network was measured twice (technical duplicate).

### 2.3. Effect of culture format on expression of fibrotic marker genes

Next, we explored how the culture format affected mRNA transcript levels of genes relevant to pathophysiological processes in endometriotic stroma and epithelium. When studying fibrosis, the biomechanical microenvironment is a highly relevant disease characteristic, as cellular fibrosis is in part regulated by biomechanical cues arising from the heterogeneous fiber network of the ECM.[37,38,53,71] Modelling this 3D biomechanical environment is therefore desirable when studying fibrosis associated with endometriosis and testing drug candidates. For T-HESCs, we chose to assess the mRNA levels of the following genes: *(i) COL1A1*, which codes for Collagen I α1, an extracellular matrix protein, *(ii) ACTA2*, which codes for α-Smooth muscle actin (α-SMA), a marker of fibroblast-to-myofibroblast transdifferentiation, *(iii) FN1*, which codes for Fibronectin-1, a protein involved in cell adhesion to the ECM which was shown to be elevated in patients with endometriosis compared to healthy patients,[72–74] and *(iv) SMAD3*, which codes for the Smad3 protein, a signal transduction molecule in the canonical TGFβ signaling pathway.[31,32,40,75–77] While a transcript-level upregulation of *COL1A1*, *ACTA2* and *FN1* indicated a pro-fibrotic state, *SMAD3* was shown to be downregulated at the mRNA level upon exposure to TGF-β1 in T-HESCs.[78] For 12Z cells, we chose to study the following genes: *(i) SNAI1*, which codes for the Snail transcription factor, *(ii) SNAI2*, which codes for the Slug transcription factor, both of which repress E-cadherin to regulate epithelial-to-mesenchymal transition, and were identified as being the transcription factors most strongly affected by TGF-β1-induced epithelial-to-mesenchymal transition in 12Z cells.[29] Further, we studied the genes *(iii) CDH1*, which codes for E-cadherin, a marker of epithelial cells involved in the formation of adherens junctions that characterize intact epithelium, and *(iv) CDH2*, which codes for N-cadherin, a marker of mesenchymal cells that functions as a promoter of cellular motility. A pro-fibrotic state in epithelial 12Z cells was characterized by an upregulation of *SNAI1*, *SNAI2* and *CDH2* and a downregulation of *CDH1* at the mRNA level.

Cells of each type, stromal T-HESC and epithelial 12Z, were harvested from monolayers and brought into suspension. This suspension of “mother cells” was used to seed FN-silk networks, spheroids, and monolayers, separately for each cell type (Figure 6A). Cells in FN-silk networks, spheroids and monolayers were cultured for 3 days in complete growth medium and subsequently for 1 day in serum-free medium. The mRNA levels of cells from FN-silk networks and spheroids were measured by qPCR and normalized to mRNA levels of cells cultured in monolayers (Figure 6A). For stromal T-HESCs, culture in FN-silk network and spheroid formats had similar effects, with mRNA levels differing by magnitudes of factor 2–3 compared to monolayers (Figure 6B). COL1A1 mRNA levels were upregulated by culture in 3D compared to monolayers, especially in spheroids, which may be attributable to more cell-cell interactions (Figure 6B). ACTA2 mRNA levels were downregulated in 3D culture compared to monolayers, suggesting that cells were in a less fibrotic state in FN-silk networks compared to monolayers (Figure 6B). ACTA2 mRNA levels may be downregulated by culture in 3D relative to monolayers, because in a monolayer format cells grow on a tissue culture-treated polystyrene surface, which is relatively stiff; while the FN-silk network as a lower, more physiological stiffness and spheroids have no surface other than other cells’ surfaces for cell adhesion. Being cultured on a stiffer mechanical microenvironment has been reported to promote fibroblast-to-myofibroblast transdifferentiation.[38,45] FN-1 mRNA, on the other hand, was upregulated in 3D culture compared to monolayers, as was SMAD3 mRNA (Figure 6B). While SMAD3 mRNA upregulation is consistent with anti-fibrotic effects in T-HESCs,[78] FN-1 mRNA upregulation is attributable to the formation of cell-cell contacts, as FN-1 mRNA was upregulated to similar magnitudes for cells in FN-silk networks and spheroids. Collagen I α1 and α-SMA proteins were expressed ubiquitously by cells in FN-silk networks (Figure 6C and D, respectively). The immunostaining provides spatial context, demonstrating the distribution and localization of these key proteins within the FN-silk network, visualizing their role in cell-matrix interactions.

**Figure 6.**
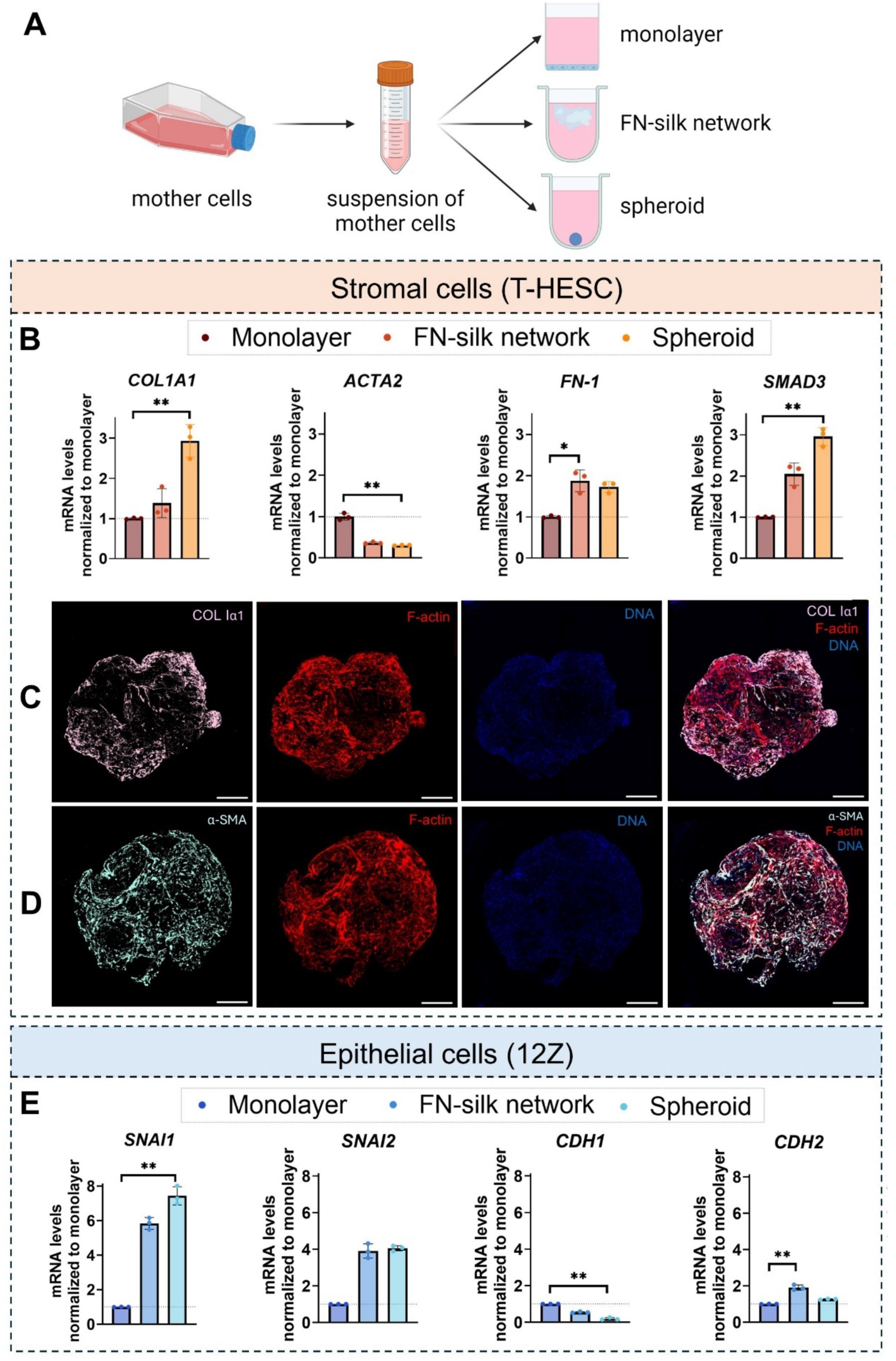
Effect of culture format on mRNA levels of transcripts involved in fibrotic pathophysiology of endometriotic stroma (T-HESC) and epithelium (12Z). **A** Schematic illustration of the seeding of different culture formats. **B** mRNA levels of *COL1A1, ACTA2, FN-1* and *SMAD3* in stromal T-HESCs grown in monolayers, FN-silk networks or spheroids were measured by RT-qPCR and normalized to mRNA levels of the reference gene *GAPDH.* The values are displayed relative to the corresponding transcript levels in the monolayer cultures (mean ± SD from N=3 independent experiments). Immunofluorescent staining for **C** Collagen I α1 and **D** α-Smooth muscle actin (α-SMA) of stromal cells in FN-silk networks. Single channel images of protein of interest, DNA (DAPI, blue) and actin filaments of the cytoskeleton (F-actin, phalloidin 594, red), and overlays. Scale bars indicate 500 µm. **E** mRNA levels of *SNAI1, SNAI2, CDH1* and *CDH2* of epithelial 12Z cells grown in monolayers, FN-silk networks or spheroids were measured by RT-qPCR and normalized to mRNA levels of the reference gene *GAPDH.* The values are displayed relative to the corresponding transcript levels in the monolayer cultures (mean ± SD from N=3 independent experiments). Statistical testing was performed using the Kruskal-Wallis test with Dunn’s multiple comparisons, and significance values were chosen as *p<0.1; **p<0.05.

For epithelial 12Z cells, mRNA levels were affected similarly in both 3D culture formats relative to monolayers (Figure 6E). However, for these cells, culture in 3D upregulated mRNA levels of transcripts involved in fibrosis. SNAI1 and SNAI2 mRNA were both upregulated in 3D cultures relative to monolayers, which corresponded with a downregulation of CDH1 mRNA relative to monolayers (Figure 6E). Concomitantly, CDH2 mRNA was upregulated in both 3D formats compared to monolayers, albeit by a small magnitude (Figure 6E). This suggests that culture in 3D appears to favor a mesenchymal-type character of epithelial 12Z cells at the mRNA level. This observation contrasts with results from Stejskalová *et al.*, who observed an upregulation of CDH1 mRNA and a downregulation of CDH2 mRNA in 12Z spheroids relative to monolayers, albeit using β-actin as a reference gene.[56] We would expect culture in 3D to increase the number of cell-cell contacts and hence downregulate epithelial-to-mesenchymal transition, however, the data presented in Figure 6E shows the opposite effect. Epithelial-to-mesenchymal transition can be caused by hypoxia, although this is unlikely in FN-silk networks, as they are thin enough (100–150 μm) to allow sufficient oxygenation of cells in the network. It may be that different mechanisms are causing epithelial-to-mesenchymal transition in each 3D system, such as hypoxia in spheroids and increased cell migration as cells proliferate in the FN-silk networks.

### 2.4. Induction of fibrosis by TGF-β1

Even though stromal T-HESCs and epithelial 12Z cells were originally isolated from patients with leiomyoma and endometriosis, respectively, and therefore already show a degree of fibrosis, fibrosis can be induced further by treating cells with TGF-β1.[17,29,41,78–81] First, we sought to characterize the effect of TGF-β1 treatment on cell metabolic activity to identify a suitable concentration to treat cells with. Figure 7A and B show the effect of TGF-β1 concentration on the viability of stromal T-HESC and epithelial 12Z cell monolayers, respectively. For T-HESCs, only 20 ng mL^-1^ TGF-β1, the highest concentration tested, reduced the viability compared to vehicle-treated cells. Other types of myofibroblasts have been observed to have higher proliferation upon exposure to TGF-β1;[82] however, this does not appear to be the case for stromal T-HESCs. For epithelial 12Z cells, TGF-β1 concentrations at 5 ng mL^-1^ and higher caused an increase in viability compared to vehicle-treated cells. This is consistent with an increase in proliferative/migratory behavior of 12Z cells observed during epithelial-to-mesenchymal transition induced by TGF-β1.[31] Figure 7C and D show that TGF-β1 treatment at 10 ng mL^-1^ for 24 h had no effect on the metabolic activity of stromal T-HESC and epithelial 12Z cells cultured in FN-silk network and spheroid formats. Thus, we proceeded with treating cells with 10 ng mL^-1^ TGF-β1.

**Figure 7.**
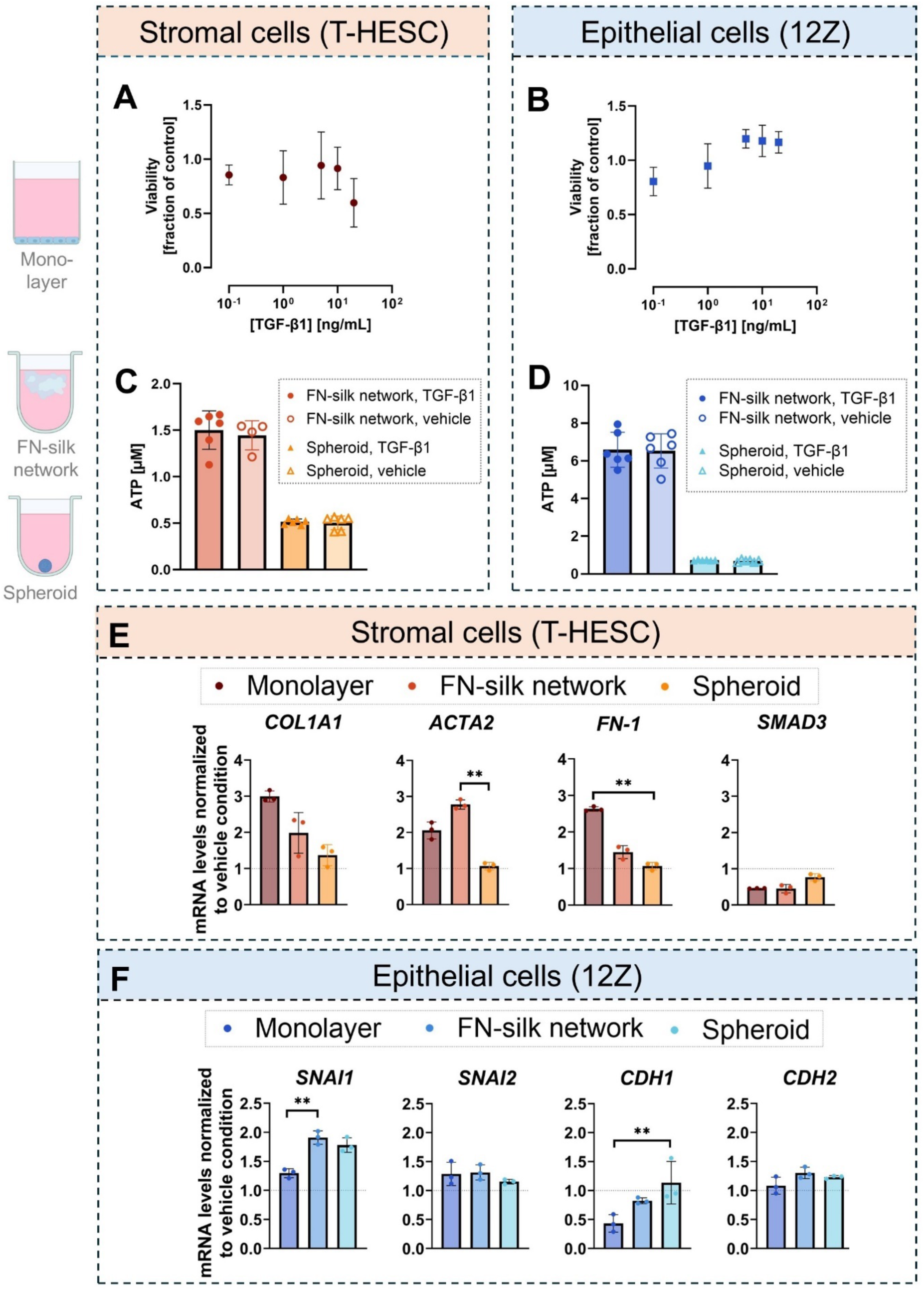
Effect of TGF-β1 concentration on the metabolic activity of **A** stromal T-HESCs in monolayers, **B** epithelial 12Z cells in monolayers (mean ± SD from N=4 independent experiments with n=3 technical replicates each), and effect of TGF-β1 (10 ng mL^-1^, 24 h) and vehicle treatment on **C** stromal T-HESCs in FN-silk networks and spheroids, and **D** epithelial 12Z cells in FN-silk networks and spheroids (mean ± SD from N=1 independent experiment with n=4−6 technical replicates). Effect of TGF-β1 on mRNA levels of transcripts involved in fibrosis in **E** stromal T-HESC and **F** epithelial 12Z cells cultured in monolayers, FN-silk networks and spheroids. mRNA levels of the transcript of interest were measured by qPCR and normalized to mean mRNA levels of the vehicle-treated unit of each respective format using the reference gene *GAPDH* (mean ± SD from N=3 independent experiments). Statistical testing was performed using the Kruskal-Wallis test with Dunn’s multiple comparisons, and significance values were chosen as *p<0.1; **p<0.05.

This concentration of TGF-β1 was not only chosen based on our observations on metabolic activity, but also because a concentration of 10 ng mL^-1^ TGF-β1 lies in the physiological range of peritoneal fluid.[32,79,83–87]

Next, we verified whether TGF-β1 treatment induced a response at the mRNA level indicative of fibrotic behavior of cells, evidencing their physiological behavior. TGF-β1 upregulated COL1A1 and ACTA2 mRNA levels for cells in monolayers and FN-silk networks, but not in spheroids (Figure 7E). The behavior of stromal T-HESCs in monolayers agrees with results described previously for primary human endometrial stromal cells for COL1A1, ACTA2, FN-1 and SMAD3 mRNA level fold changes.[78] This indicates that stromal T-HESCs are a suitable cell line to model fibrotic pathophysiology, as their behavior mimics physiological behavior of primary human endometrial stromal cells at the mRNA level. Even though stromal T-HESCs were inherently in a myofibroblast-like state, as they were isolated from a donor with leiomyoma,[88] TGF-β1 further augmented the myofibroblast-like character of these cells. The magnitude of COL1A1 and ACTA2 mRNA upregulation was greater for cells in monolayers (upregulation by 192–217%) than in FN-silk networks (upregulation by 34 –134%), although the difference was not statistically significant (Figure 7E). This indicates that culturing stromal T-HESCs in an FN-silk network format made them less responsive to TGF-β1 than monolayers; which may be because the FN-silk matrix has a lower, more tissue-relevant stiffness and provides less mechanical tension. Lower responsiveness of T-HESCs in 3D to TGF-β1 may also be due to differences in cell-cell adhesion and integrin signaling. Studying fibrosis in monolayers may cause us to overestimate effect magnitudes compared to conditions mimicking the *in vivo* environment more closely, although the trends are similar.

Figure 7E also shows that FN-1 mRNA was upregulated to a greater magnitude for cells in monolayers (factor 160–170%) than in FN-silk networks (factor 30–60%), although this difference was not large enough to be statistically significant. This may be because the mRNA levels of FN-1 were already upregulated by culture in an FN-silk network format relative to a monolayer (Figure 6B), as more cell-cell and cell-ECM contacts are formed, and TGF-β1 did not further upregulate mRNA levels of Fn-1. SMAD3 mRNA levels, on the other hand, were downregulated by TGF-β1 treatment in all three formats, to a greater extent in monolayers (downregulation by 50%) and FN-silk networks (downregulation by 50–70%) than in spheroids (downregulation by 9–29%) (Figure 7E). This downregulation of SMAD3 mRNA levels is consistent with results from monolayers of primary human endometrial stromal cells, where TGF-β1 treatment induced downregulation of SMAD3 mRNA levels by ∼60%.[78] In our results, SMAD3 mRNA levels were inversely correlated with COL1A1 and ACTA2 mRNA levels relative to the vehicle-treated control.

Stromal T-HESC spheroids showed little response to TGF-β1 at the mRNA level, showing no upregulation in COL1A1, ACTA2, and FN-1 mRNA levels compared to vehicle-treated spheroids (Figure 7E). This suggests that either stromal T-HESCs in spheroids were in a state where they did not respond to cytokines (either dead, or in a dormant state), or that the spheroid architecture was so dense that TGF-β1 was unable to penetrate to cells inside the spheroid and only acted on cells in the outermost layer of the spheroid. Either way, our results suggest that spheroids are unsuitable to model fibrosis in endometrial stroma in 3D.

For epithelial 12Z cells, TGF-β1 upregulated SNAI1 mRNA levels in all three formats (Figure 7F). The magnitude of SNAI1 mRNA upregulation was greater for cells in FN-silk networks (upregulation by 80–100%) and spheroids (upregulation by 70–90%) than in monolayers (upregulation by 20–40%), although the difference was only statistically significant between monolayers and FN-silk networks (Figure 7F). Epithelial 12Z cells cultured in all three formats showed a slight upregulation of SNAI2 mRNA levels, however, the magnitude was small, ranging from 10-50%, suggesting that TGF-β1 acted primarily *via* SNAI1 and to a lesser extent *via* SNAI2 at the mRNA level in epithelial 12Z cells (Figure 7F). This is consistent with observations from Ma *et al.* on epithelial 12Z cells, who also observed a greater magnitude of upregulation of SNAI1 mRNA levels than SNAI2 mRNA levels upon TGF-β1 treatment.[29] Epithelial 12Z cells in spheroids responded to TGF-β1 exposure at the mRNA level (Figure 7F), hinting that 12Z cells in spheroids were more responsive than T-HESC spheroids, and that epithelial 12Z spheroid architecture allowed TGF-β1 to penetrate to the center of the spheroid. Our observations suggest that for epithelial 12Z cells, a loss of epithelial character was not coupled to a gain in mesenchymal character.

Despite epithelial 12Z monolayers showing the smallest upregulation in SNAI1 mRNA, CDH1 mRNA was downregulated by the largest magnitude for cells in monolayers (downregulation by 50–70%) compared to FN-silk networks (downregulation by 10–23%) and spheroids (no change) (Figure 7F). This shows that culture in the monolayer caused TGF-β1 to have a greater effect on loss of epithelial character than culture in the FN-silk networks. Compared to monolayers, CDH1 mRNA was already downregulated in both 3D culture formats (Figure 6E), and TGF-β1 treatment did not further downregulate CDH1 mRNA in 3D culture (Figure 7F). This may be because the FN-silk matrix had a lower stiffness than a tissue-culture treated flat polystyrene surface. Unexpectedly, we observed no change in CDH2 mRNA levels upon TGF-β1 treatment in any of the three culture formats (Figure 7F). For a complete epithelial-to-mesenchymal transition, we would expect CDH2 mRNA levels up be upregulated, as *CDH2* codes for N-cadherin, a mesenchymal marker. Taken together, these results suggest that TGF-β1 treatment caused a loss in epithelial character of the 12Z cells, to the greatest extent in monolayers, but little gain of mesenchymal character. In contrast, Ma *et al.* observed an upregulation of CDH2 in 12Z cell monolayers by a factor of ∼50% upon TGF-β1 treatment.[29] Our observations are consistent with the partial epithelial-to-mesenchymal transition process observed in primary human endometriotic epithelial cells[42] and suggested by Konrad *et al.*[30]

The effect of TGF-β1 observed in monolayer cultures was reproduced in FN-silk networks for stromal T-HESCs, where it induced fibroblast-to-myofibroblast transdifferentiation, but not in spheroids. In epithelial 12Z cells, TGF-β1 affected epithelial-to-mesenchymal transition across all formats, indicated by a loss of epithelial character with minimal mesenchymal gain.

### 2.5. Testing pirfenidone, an antifibrotic drug candidate with the potential to be repurposed for endometriosis treatment

Exploring the similarities between endometriosis and other fibrotic diseases, such as idiopathic pulmonary fibrosis, could provide valuable insights for the development of pharmacological treatments for endometriosis.[10] Pirfenidone is a small molecule drug indicated for idiopathic pulmonary fibrosis, a disease that shares common mechanisms with endometriosis. Pirfenidone acts by regulating the Wnt/β-catenin and TGF-β1/Smad2/3 signaling pathways.[89,90] In a randomized, double-blind, prospective clinical trial with 210 patients undergoing laparoscopic surgery for endometriosis, it was found that 1800 mg of oral pirfenidone administered daily for six months post-surgery led to a reduction in fibrotic adhesions compared to the placebo group.[62] We chose to test the antifibrotic activity of pirfenidone on *in vitro* models of fibrosis in endometriosis developed here to evaluate their suitability as screening tools for antifibrotic drug candidates.

Part of pirfenidone’s mechanism of action is that it inhibits the proliferation of myofibroblasts. However, we were interested in pirfenidone’s effect on the expression of genes involved in fibrotic pathophysiology, rather than cell proliferation. As such, we sought to identify a pirfenidone concentration that was high enough to be likely to affect mRNA levels, but low enough not to inhibit cell proliferation. To this end, we assessed the effect of the drug concentration on the viability of stromal T-HESCs and epithelial 12Z cells in monolayers. For both cell types, viability remained close to untreated cells at pirfenidone concentrations of up to 1 mg mL^-1^, while concentrations above 1 mg mL^-1^ reduced viability (Supporting Information, **Figure S.8**). Therefore, we chose to treat cells with pirfenidone at a concentration of 0.8 mg mL^-1^ for drug testing experiments.

We investigated the effect of pirfenidone on mRNAs involved in fibrosis in stromal T-HESCs and epithelial 12Z cells cultured in monolayers, FN-silk networks, and spheroids (Figure 8A). Each cell type/format combination was pre-treated with TGF-β1 or vehicle solution for 24 h and subsequently treated with either pirfenidone in serum-free medium or serum-free medium only for 24 h (Figure 8B). Stromal T-HESCs were treated with 0.82 ± 0.03 mg mL^-1^ pirfenidone and epithelial 12Z cells with 0.85 ± 0.04 mg mL^-1^ pirfenidone (mean ± SD from N=3 independent experiments). We additionally chose to pre-treat cells with TGF-β1 to assess whether pirfenidone can reverse fibrosis caused by the pre-treatment, making the model more relevant to *in vivo* conditions in which fibrosis is already present in endometriotic tissue before a drug is given.

**Figure 8.**
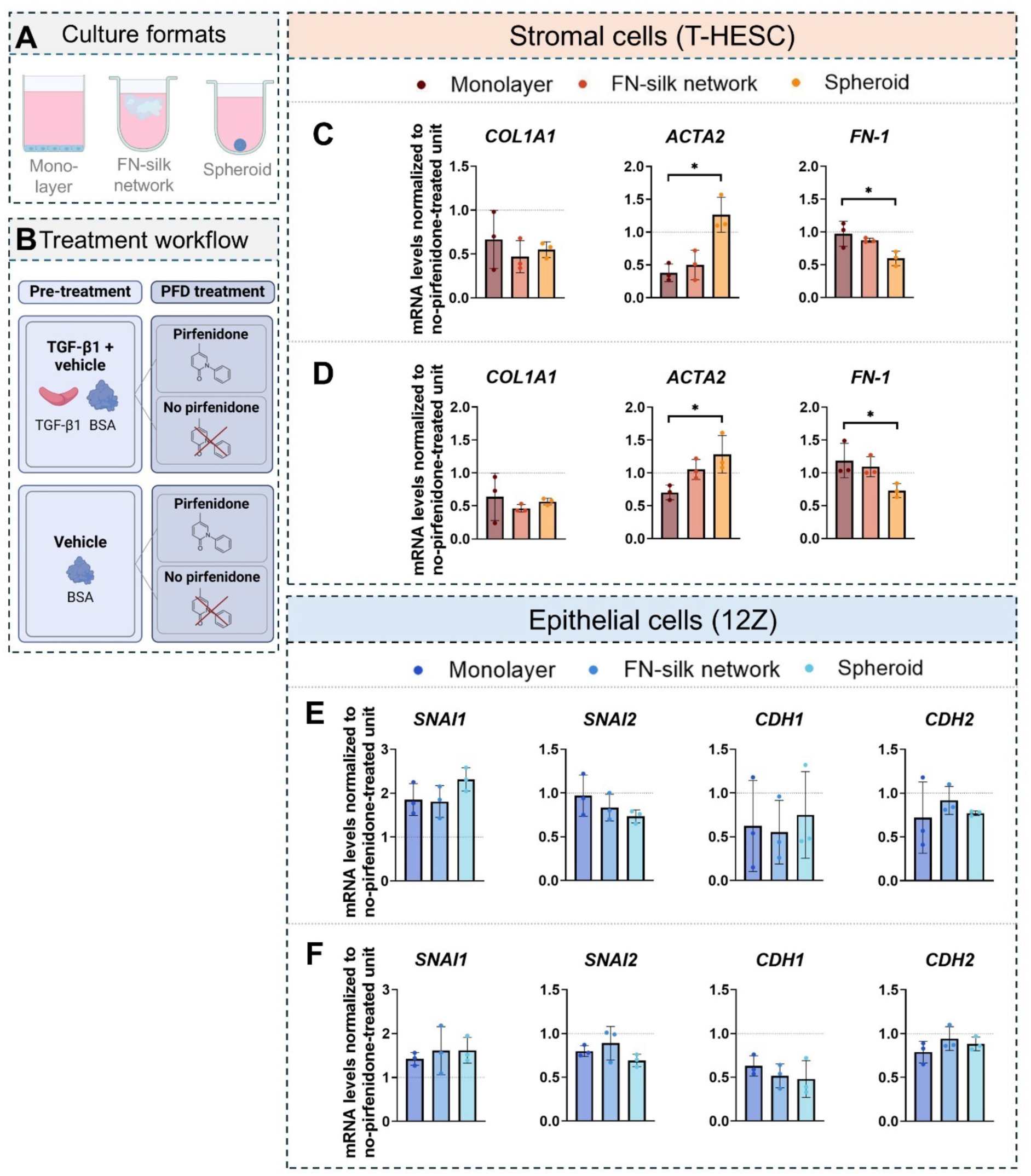
Treatment of stromal T-HESC and epithelial 12Z cells with pirfenidone. **A** Schematic illustration of 2D (monolayer) and 3D (FN-silk network, spheroid) culture formats. **B** Treatment workflow, illustrating pre-treatment with TGF-β1 or vehicle, followed by treatment with pirfenidone or no pirfenidone in serum-free medium for each respective pre-treatment. This treatment workflow was applied to cells in each culture format. Effect of pirfenidone on mRNA levels of transcripts involved in fibrosis, measured using RT-qPCR, normalized to the mean of no-pirfenidone treated cells cultured in each respective format using the reference gene *GAPDH*. **C** Stromal T-HESCs pre-treated with TGF-β1, **D** stromal T-HESCs pre-treated with vehicle, **E** epithelial 12Z cells pre-treated with TGF-β1, **F** epithelial 12Z cells pre-treated with vehicle (mean ± SD from N=3 independent experiments). Statistical testing was performed using the Kruskal-Wallis test with Dunn’s multiple comparisons, and significance values were chosen as *p<0.1; **p<0.05.

Figure 8C shows the effect of pirfenidone on COL1A1, ACTA2 and FN-1 mRNA levels for stromal T-HESCs pre-treated with TGF-β1. Pirfenidone downregulated COL1A1 mRNA levels in stromal T-HESCs in monolayers (reduction by 2–68%), FN-silk networks (reduction by 32– 66%), and spheroids (reduction by 38–55%). To our surprise, stromal T-HESC spheroids did appear to respond to pirfenidone treatment in terms of COL1A1, suggesting that the small molecule was able to diffuse into the core of the spheroid. Pirfenidone downregulated ACTA2 mRNA levels to similar magnitudes for stromal cells in monolayers (reduction by 50–70%) and FN-silk networks (reduction by 29–70%) (Figure 8C). For vehicle pre-treated cells, pirfenidone hardly downregulated ACTA2 mRNA in monolayers (reduction by 25–38%), and not at all in FN-silk networks, indicating that pirfenidone reversed upregulation of fibrosis-related mRNA transcripts induced specifically by TGF-β1 (Figure 8D). T-HESC spheroids, however, did not react to pirfenidone treatment in terms of ACTA2 mRNA levels (Figure 8C and D), suggesting that stromal T-HESC spheroids may be less well suited for testing antifibrotic drugs. Further, Figure 8C and D shows that pirfenidone had no effect on FN-1 mRNA levels of stromal T-HESCs neither in monolayers nor in FN-silk networks, indicating that pirfenidone did not reverse TGF-β1-induced FN-1 mRNA upregulation (Figure 8C and D). Surprisingly, T-HESC spheroids treated with pirfenidone showed a small downregulation of FN-1 mRNA levels in both TGF-β1 and vehicle pre-treatments (reduction by 29–50%). Together with our observations that stromal T-HESC spheroids were unresponsive to TGF-β1 at the FN-1 mRNA level (Figure 7E), this suggests that likely the culture in serum-free medium caused in reduction in FN-1 mRNA levels in stromal T-HESC spheroids.

Results shown in Figure 8C and D validated our screening platform: cellular fibrosis was reversed at the mRNA level when stromal cells were exposed to a known antifibrotic compound, pirfenidone. Pirfenidone downregulated mRNA levels of COL1A1 and ACTA2, hinting that this drug may have the potential to prevent ECM protein deposition and possibly reverse TGF-β1-induced fibroblast-to-myofibroblast transdifferentiation in endometriotic stroma. Pirfenidone’s anti-fibrotic activity at the mRNA level was indicated similarly for stromal T-HESCs in monolayers and FN-silk networks, and to a lesser extent in spheroids.

Although pirfenidone’s efficacy has been mainly attributed to its action on fibroblasts,[91] we were interested in studying the effect of pirfenidone on epithelial cells, which form a substantial fraction of cells in the endometrium and endometriotic lesions.[92] Figures 8E **and F** show that 12Z cells generally behaved similarly in all three culture formats in terms of SNAI1, SNAI2, CDH1 and CDH2 mRNA levels, relative to the no-pirfenidone treated epithelial 12Z cells in each respective culture format. In TGF-β1 pre-treated 12Z cells, pirfenidone treatment upregulated SNAI1 mRNA levels by a factor of 40–160% and did not affect SNAI2 mRNA levels (Figure 8E). The trend of SNAI1 mRNA upregulation by pirfenidone was similar although slightly smaller in vehicle pre-treated 12Z cells across all culture formats (Figure 8F). CDH1 transcription was unchanged in TGF-β1 pre-treated 12Z cells across all culture formats (Figure 8E) and downregulated by a factor of 20–71% in control pre-treatments (Figures 8F), suggesting that pirfenidone promoted a loss of epithelial character by upregulating SNAI1 mRNA levels. However, CDH2 transcription remained largely unaffected by pirfenidone treatment.

Taken together, these mRNA-level results from 12Z cells across all culture formats provide tentative evidence that pirfenidone facilitated epithelial-to-mesenchymal transition in 12Z cells independent of pre-treatment, albeit to a small magnitude. Together with evidence of pirfenidone’s effect on other types of epithelial cells,[93–97] our results suggest that this drug’s primary anti-fibrotic mechanism of action stems from its effect on fibroblasts by reversing fibroblast-to-myofibroblast transdifferentiation, rather than its effect on epithelial cells.

## 3. Conclusion and outlook

In this study, we have developed 3D culture models of fibrosis in endometriosis and used these models to evaluate a candidate antifibrotic drug with the potential to be repurposed for endometriosis. We demonstrated that FN-silk networks, developed by Johansson, Widhe *et al.*[46] and Collodet *et al.*[47] and adapted slightly here, were a suitable matrix to culture endometrial stromal cells (T-HESCs) in 3D, while spheroids were less suitable in terms of their ability to mimic the fibrotic character of stromal cells found in endometriosis. In addition to being fully defined, non-animal origin, and readily available, FN-silk as a scaffold material was shown to be advantageous due to its tissue-mimicking stiffness. We showed that stromal cells cultured in FN-silk behaved physiologically in terms of cell growth and mRNA-level response to the profibrotic cytokine TGF-β1, while culture of stromal T-HESCs in a spheroid format did not enable this physiological behavior to be modelled as comprehensively. For epithelial cells, FN-silk and spheroid formats partially modelled fibrotic processes in 3D culture, as probed at the mRNA level, indicating that both formats were suitable to model fibrosis in epithelial 12Z cell cultured in 3D.

Testing pirfenidone on these *in vitro* models provided evidence for the drug’s anti-fibrotic effects on endometriotic stroma by downregulating mRNA levels of transcripts implicated in fibroblast-to-myofibroblast transdifferentiation. This demonstrated that the stromal cell FN-silk network 3D culture format is an effective drug screening tool for evaluating antifibrotic candidates. However, pirfenidone upregulated epithelial-to-mesenchymal transition-promoting mRNA levels in epithelial 12Z cells in 3D, albeit to a small magnitude, highlighting the drug’s distinct effects on stromal and epithelial components of endometriotic lesions. These observations underscore the utility of *in vitro* systems to closely characterize the behavior of different cell types in response to drug treatment. In this study, we used qPCR to reliably and quantitatively compare culture formats in terms of changes to their gene expression patterns in response to TGF-β1 and pirfenidone treatment. For future applications of these models in drug testing, additional protein-level methods should be considered to provide a more comprehensive understanding of the candidate drug’s effects. In terms of developing the model further, future work should focus on bringing stromal and epithelial cells into co-culture on an FN-silk network, as this co-culture model may allow the precise orchestration of TGF-β signaling between stromal and epithelial cells[92] to be interrogated *in vitro*.

By continuing to develop and refine these sustainable *in vitro* models, we can enhance pharmaceutical research on endometriosis while aligning with the UN SDGs, ultimately contributing to both innovative treatments for women’s health and the promotion of environmentally responsible research practices. Our work demonstrates that an SDG-guided framework can be effectively applied at various stages of pharmaceutical research and development, from project selection to methodology. By focusing on women’s health and minimizing animal experiments, we ensure that our research contributes to sustainable practices that are both ethically sound and scientifically rigorous. This work paves the way for more efficient, responsible, and potentially meaningful approaches to tackling complex diseases like endometriosis, underscoring the importance of integrating sustainable development goals into the heart of pharmaceutical research.

## 4. Materials and methods

### 4.1. Cell culture

Immortalized human endometrial stromal cells (T-HESC cell line,[88] Cellosaurus accession no. CVCLC464, female sex) were purchased from Applied Biological Materials, Canada (Ref T0533). T-HESCs were cultured at 37 °C in a humidified atmosphere containing CO2 (5%). The complete growth medium composition was DMEM/F12 without phenol red (Gibco™ Ref 21041025, Paisley, UK), charcoal-stripped fetal bovine serum (10 v/v%, Gibco™ Ref 12676029, Grand Island, NY, USA), ITS+ Premix Universal Culture Supplement (1 v/v%, Corning™ Ref 354352, Bedford, MA, USA [final concentrations in medium: human recombinant insulin (5 μg mL^-1^); transferrin (5 μg mL^-1^); selenous acid (5 ng mL^-1^)]) and penicillin-streptomycin (1v/v%, Gibco™ Ref 15140122, Grand Island, NY, USA). Subcultivation of monolayers was performed at 70–80% confluence by detachment using TrypLE (Gibco™ Ref 12604013, Paisley, UK). For experiments, cells from passage numbers 5–18 were used, where passage number 0 was defined as the delivered vial. Genomic profiling was done at passage 8 using highly polymorphic short tandem repeat loci (STRs). STR loci were amplified using the PowerPlex^®^ 16 HS System (Promega), fragment analysis was done on an ABI3730xl (Life Technologies) and the resulting data were analyzed using the GeneMarker HID software (Softgenetics). A 100% match to the DNA profile of the T-HESC line was found. Full STR profile results are shown in the Supporting Information (**Table S.1, Figure S.9**).

Immortalized human epithelial endometriotic cells (12Z cell line,[98] Cellosaurus accession no. CVCL0Q73, female sex) were purchased from Applied Biological Materials, Canada (Ref T0764). 12Z cells were cultured at 37 °C in a humidified atmosphere containing CO2 (5%). The complete growth medium composition was Dulbecco’s Modified Eagle Medium (DMEM) with glucose (4.5 g L^-1^), without sodium pyruvate (Gibco™ Ref 11960044, Paisley, UK), fetal bovine serum (10 v/v%, Merck EMD Millipore, Ref ES-009-B, Burling ton, MA, USA), L-glutamine (200 mM, Gibco™ 25030024, Paisley, UK) and penicillin-streptomycin (1 v/v%, Gibco™ Ref 15140122, Grand Island, NY, USA). Subcultivation of monolayers was performed at 70–90% confluence by detachment using TrypLE (Gibco™ Ref 12604013, Paisley, UK). For experiments, cells from passage numbers 9-22 were used, where passage number 0 was defined as the delivered vial. Genomic profiling was done at passage 14 and a 96% match to the STR profile of the 12Z cell line was found. Differences were found in the D8S1179 and FGA loci (**Table S.2, Figure S.10**). Cultures were routinely tested for mycoplasma (MycoStrip Mycoplasma Detection Kit, InvivoGen, San Diego, CA, USA) and no mycoplasma contamination was detected.

### 4.2. 3D culture

#### 4.2.1. FN-silk networks

FN-silk networks were prepared as described previously[46,47] with adaptations, as shown in Figure 2A. Cells were harvested and brought into suspension in complete growth medium. FN-silk in PBS (Spiber Technologies AB, Stockholm, Sweden) was freshly thawed and mixed with the cell suspension at a 2:1 volumetric ratio to make a master mix containing liquid FN-silk protein and suspended cells. The final concentration of FN-silk in the master mix was 2.2 mg mL^-1^ and the cell concentration in the master mix was 1×10^6^ cells mL^-1^ for T-HESC and 2×10^6^ cells mL^-1^ for 12Z cells, such that the number of cells per FN-silk network was 10,000 cells for T-HESC and 20,000 cells for 12Z. A droplet of master mix (10 μL) was placed at the bottom of each well in a U-shaped-bottom ultralow attachment plate (Nunclon Sphera-Treated U-Bottom Microplate, Thermo Fisher Scientific Ref 174929, Japan). A multichannel pipette set to 28 μL was used to rapidly pipette air into the mastermix droplet 30 times, to allow the silk protein to assemble into β-sheets at the air-liquid interface and thereby form a foam. The foam was incubated in a humidified incubator at 37 °C for 12 min. Each foam was suspended in complete growth medium (180 μL) and, using a μ-spoon, transferred to a new well containing fresh complete growth medium (180 μL). The FN-silk foam transformed into a network as the sheets around the bubbles burst. Cells in FN-silk networks were cultured for up to 7 days and medium was changed every 2−3 days. At day 2, all air bubbles had disappeared and FN-silk networks were submerged below the surface of the medium in the center of the well. Integration of cells in FN-silk networks was measured by counting the cells left in the medium after transferring the FN-silk network to a well containing fresh medium. Cells were stained with a Trypan blue solution (0.4%, Merck Ref T8154, Switzerland) and counted using a Neubauer plate.

#### 4.2.2. Spheroids

Spheroids were prepared by seeding a cell suspension into U-shaped-bottom ultralow attachment plate (Nunclon Sphera-Treated U-Bottom Microplate, Thermo Fisher Scientific Ref 174929, Japan) and allowing cells to spontaneously aggregate into spheroids, as described previously[63] and illustrated in Figure 2B. The number of cells seeded per well was equivalent to the FN-silk networks. Spheroids were cultured in 100 μL medium per well for up to 7 days and medium was changed every 2−3 days.

### 4.3. Metabolic activity

#### 4.3.1. Monolayers

The viability of cells cultured in monolayers was measured using the Cell Counting Kit-8 (CCK-8, Merck Ref 96992, Switzerland). The CCK-8 assay was performed according to manufacturer’s instructions and as described in[99]. The absorbance was measured at *λ* = 450 nm using a plate reader (Tecan Infinite M Nano, Switzerland), with *λ* = 650 nm as the reference wavelength. To calculate the viability, Equation 1 was used, where *OD*_*sample*_ and *OD*_*control*_ are the optical densities of treated and control cell monolayers, respectively.

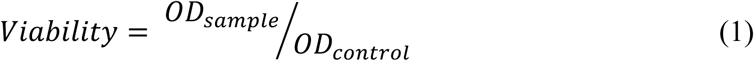

#### 4.3.2. 3D systems

The metabolic activity of cells cultured in 3D formats was measured using the CellTiter Glo^®^ Cell 3D Viability Assay (Promega Ref G9681, Madison, WI, USA). The CellTiter Glo 3D Viability Assay was performed according to the manufacturer’s instructions with minor modifications. All materials were equilibrated to 24 °C and all steps performed in low-light conditions. Each 3D cell unit was transferred to a well of a white plate (pureGrade S polystyrene flat-bottom 96 well plate, Brand Ref 781665, Wertheim, Germany) containing complete growth medium (80 μL) using a μ-spoon for FN-silk networks and wide-orifice pipette tips (Finntip Wide Orifice Pipette Tips, Thermo Fisher Scientific Ref 9405163) for spheroids. Cell systems were equilibrated to 24 °C for 25 min in the dark. CellTiter Glo 3D Viability reagent (80 μL) was added to each well and the contents were mixed vigorously for 5 min to induce cell lysis. The plate was incubated for 25 min in the dark and luminescence measured using a plate reader (Tecan Infinite 200 PRO F Plex/M Nano+, Switzerland). The CellTiter Glo Cell 3D Viability Assay was validated by comparison to a manual count of cells (Supporting Information **Figure S.4**).

In addition to metabolic assays, the viability of cells in 3D systems was assessed visually using a live/dead staining kit (LIVE/DEAD Viability/Cytotoxicity Assay Kit, Invitrogen Thermo Fisher Scientific Ref L32250, Eugene, OR, USA). The live/dead assay was performed according to the manufacturer’s instructions. Briefly, a working solution with composition SYTOX Deep Red Nucleic Acid Stain (0.25 μM) and calcein AM (2 μM) in complete growth medium was prepared. Cells were incubated in the working solution at 37 °C, 5% CO2 for 30 min in the dark. FN-silk networks and spheroids were visualized directly in the well using a wide-field microscope (Kinetix Ti2, Nikon, Japan), with the GFP filter set for calcein-stained cells and the deep red/Cy5 filter set for SYTOX Deep Red-stained cells.

### 4.4. Fibrosis induction

A profibrotic state was induced in both T-HESC and 12Z cells by treatment with human recombinant TGF-β1 (Stemcell Ref 78067, USA). TGF-β1 was reconstituted in hydrochloric acid (HCl, 10 mM) and diluted in a vehicle solution of bovine serum albumin (BSA, final concentration 0.1 w/v%) according to the manufacturer’s instructions. Unless otherwise stated, cells were treated with TGF-β1 at a concentration of 10 ng mL^-1^ in serum-free medium for 24 h. The corresponding composition of the vehicle, which was used as a control, was 0.0001 w/v% BSA, 1 μM HCl (final concentration in serum-free medium).

### 4.5. Pirfenidone treatment

Pirfenidone (5-methyl-1-phenylpyridin-2(1H)-one) was purchased from BLD Pharm, Germany. Pirfenidone was dissolved in serum-free medium and the concentration measured by absorbance at *λ* = 311 nm in a black/clear bottom quartz microplate using a plate reader (Tecan Infinite 200 PRO F Plex/M Nano+, Switzerland). Unless otherwise stated, T-HESCs were treated with 0.82 ± 0.03 mg mL^-1^ and 12Z cells with 0.85 ± 0.04 mg mL^-1^ pirfenidone (mean ±SD, N = 3 independent experiments) in serum-free medium for 24 h. First, cells were treated with TGF-β1 for 24 h to induce a fibrotic state, followed by treatment with pirfenidone for 24 h. Control pre-treatments were vehicle in serum-free medium and complete growth medium.

### 4.6. Immunofluorescent staining, wide field and confocal microscopy

Whole FN-silk networks were stained for EpCAM, vimentin, Collagen I α1 (COL I α1) and α-smooth muscle actin (α-SMA) using immunofluorescent staining. FN-silk networks were fixed in paraformaldehyde (4%, PFA, Thermo Fisher Scientific Ref J61899-AK) in PBS for 20 min at 24 °C and stored in PBS at 4 °C until staining. Networks were incubated in Sudan Black (SB, Thermo Fisher Scientific Ref 190160250, 0.3 w/v% in 70 v/v% ethanol, filtered through a 0.2 µm polyethersulfone membrane) for 15 min to minimize autofluorescence of the FN-silk protein. After washing with PBS three times, cells were permeabilized with Triton-X100 (0.2% in PBS, Thermo Fisher Scientific Ref A16046.AP) for 10 min. FN-silk networks were washed with Tween 20 (0.1% in PBS, Sigma-Aldrich Ref 822184) for 10 min while agitating. Cells were blocked by incubating in goat serum (1 v/v% in PBS/Tween 20 0.1%, Capricorn Scientific Ref GOA-1A) for 1 h. The primary antibody (**Table 1**) was diluted in goat serum (1 v/v% in PBS/Tween 20 0.1%) and FN-silk networks incubated with the primary antibody overnight at 4°C. FN-silk networks were washed with Tween 20 (0.1% in PBS) for 10 min while agitating. The secondary antibody (**Table 1**) was diluted in 1 v/v% goat serum (1 v/v% in PBS/Tween 20 0.1%) and FN-silk networks incubated with the secondary antibody for 2 h at 24 °C in the dark. FN-silk networks were incubated with SB for 10 min. SB was rinsed off the networks by washing with Tween 20 (0.1% in PBS) three times. Nuclei were counterstained with 4′,6-diamidino-2-phenylindole (DAPI, Carl Roth, Karlsruhe, Germany, Ref 6335.1) and F-actin was stained with Alexa Fluor 594 Phalloidin (Invitrogen™ Ref A12381, Eu-gene, OR, USA). FN-silk networks were washed with Tween 20 (0.1% in PBS) for 10 min while agitating. As a control, cells were stained with secondary antibody only (no primary antibody) to check for any unspecific binding (Supporting Information, **Figure S.11**).

**Table 1.** Antibodies used for immunofluorescent staining.

| Name | Class, isotype | Dilution | Ref no. | Supplier |
| --- | --- | --- | --- | --- |
| Primary antibodies |  |  |  |  |
| Vimentin<br>Rabbit anti-human | Monoclonal IgG | 1:500 | ab92547 | Abcam |
| EpCAM<br>Rabbit anti-human | Monoclonal IgG | 1:500 | ab223582 | Abcam |
| COL I α1<br>Rabbit anti-human | Polyclonal IgG | 1:500 | ab34710 | Abcam |
| α-SMA<br>Rabbit anti-human | Polyclonal IgG | 1:500 | PA5-19465 | Thermo Fisher Scientific |
| Secondary antibody |  |  |  |  |
| AlexaFluor 488<br>Goat anti-rabbit | Polyclonal IgG | 1:1000 | A32731 | Thermo Fisher Scientific |

For wide-field microscopy, a drop of DAKO fluorescence mounting medium (Agilent Ref S302380-2, Santa Clara, CA, USA) was placed on a glass slide, the network placed into the mounting medium using a µ-spatula, covered with a glass cover slip, and sealed around the edges using nitrocellulose (nail varnish). Networks were visualized using a Ti2 Kinetix wide-field microscope with a 10X/0.45 air objective (Nikon, Tokyo, Japan).

For confocal microscopy, DAKO mounting medium (40 µL) was placed in each well of a glass-bottom µ-well slide (15 well 3D glass bottom µ-slide, ibidi Ref 81507, Gräfelfing, Germany). Each FN-silk network was transferred to a well using a µ-spatula and visualized using a TCS SP8 DLS Light Sheet microscope in confocal mode with a 10X air objective (Leica Microsystems, Wetzlar, Germany).

### 4.7. RNA extraction, cDNA synthesis, and qPCR

#### 4.7.1. RNA extraction

Total RNA was extracted using the ReliaPrep^TM^ Cell and Tissue Miniprep System (Promega Ref Z6211, Madison, WI, USA) according to Section 5 of the manufacturer’s protocol (RNA Isolation and Purification from Cell Samples), with minor adaptations. Cells were harvested from monolayers using TrypLE. 3D systems were pooled by placing 14−16 units (FN-silk networks or spheroids) in a microcentrifuge tube with PBS. Cells were centrifuged, the supernatant removed, and the pellet frozen at −80 °C. Cells in a frozen pellet were lyzed and 3D systems broken up by pipetting up and down 100 times. The nucleic acid/cell debris mix was diluted and centrifuged at 12000×g for 5 min to pellet FN-silk and cell debris. Taking the supernatant, nucleic acids were bound to a Reliaprep MiniColumn, washed, eluted, and treated with DNase. RNA was bound to a second Reliaprep MiniColumn. After washing, RNA was eluted and quantified using spectrophotometry at λ = 260 nm (NanoDrop 2000, Thermo Fisher Scientific, USA).

#### 4.7.2. cDNA synthesis

RNA was reverse transcribed to complementary DNA (cDNA) using the GoScript^TM^ Reverse Transcriptase Random Primers kit (Promega Ref A2801, Madison, WI, USA) according to the manufacturer’s instructions.

#### 4.7.3. qPCR

cDNA levels were measured by quantitative PCR using the GoTaq^®^ qPCR kit (Promega Ref A6002, Madison, WI, USA) according to the manufacturer’s instructions under standard cycling conditions. Thermal cycling and fluorescence acquisition were performed in a Rotor-Gene Q 2Plex System (Qiagen, Germany). RT minus reactions were run in every experiment to control for DNA contamination. Relative quantification was performed using the ΔΔ*Ct* method[100] using *GAPDH* as a reference gene. Primers were synthesized by Microsynth (Balgach, Switzerland) and had amplification efficiencies of 80–98%, similar to GAPDH, over a 100-fold concentration range, allowing the ΔΔ*Ct* method to be used.[100] Primer sequences and validations are provided in the Supporting Information (**Table S.3**). *GAPDH* was found to be a suitable reference gene, as its mRNA levels were unaffected by TGF-β1 treatment (Supporting Information, **Figure S.12**).

### 4.8. Mechanical characterization

Mechanical properties of stromal T-HESC FN-silk networks were characterized on culture days 3 and 6 using a UNHT^3^ Bio Bioindenter (Anton Paar, Graz, Austria) with a cylindrical flat punch indenter (diameter 1 mm). One FN-silk network was placed on the bottom of a glass-bottom well (inner diameter 4 mm, 15 well 3D glass bottom µ-slide, ibidi Ref 81507, Gräfelfing, Germany) and PBS (100 μL) added to the well. The stiffness and elastic modulus of the FN-silk network was measured by compressing the FN-silk network with the indenter with a linear loading profile of the normal force *Fn* while recording the penetration depth Pd (Figure 5A). The loading rate was 120 μN min^-1^ up to a maximum nominal indentation load of 25 μN, followed by a pause of 5 s and unloading at a rate of 120 μN min^-1^. The penetration depth *Pd* was recorded at a rate of 50 Hz. Measurements were performed at 22 °C. As a control, a well containing only PBS and no sample was measured. The compressive elastic modulus *E* was evaluated according to Equation 1.

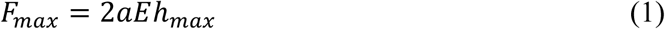

where *F*_*max*_ is the maximum applied force, *a* is the radius of the indenter shaft (here, *a* = 0.5 mm), and ℎ_*max*_ is the maximum penetration depth. Additionally, we estimated the elastic modulus *EIT* according to the Oliver & Pharr model[70] (ISO 14577) as described by Equation 2.

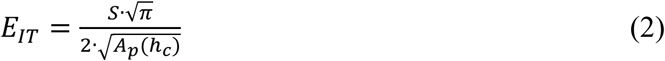

where *S* is the unloading stiffness (slope of the unloading curve at *Fmax*) and *Ap(hc)* is the projected contact area at contact depth *hc*. For a cylindrical flat punch, the projected contact area is constant (independent of the penetration depth), *A*_p_ = π*a*^2^, where *a* is the radius of the indenter shaft. The Oliver & Pharr model applies strictly to materials such as metals, glass and ceramics, as it assumes that the material is homogeneous, linearly elastic, isotropic and incompressible.[101] Here, we use the Oliver & Pharr model[70] to estimate the elastic modulus of FN-silk networks to allow a comparison to elastic moduli of other materials. However, since the FN-silk network does not satisfy all assumptions of the Oliver & Pharr model, the *EIT* values are merely estimates. Equation 2 also relies on the assumption that the stiffness of the indenter shaft is much greater than that of the sample, which was satisfied.

### 4.9. Morphological analysis

The size of cells in monolayers, of FN-silk networks and spheroids was measured by an automated method using CellPose, a generalist algorithm for cellular segmentation.[102] Code was written and executed in Python 3.10.13 in a Conda 23.11.0 environment and is available on GitHub at https://github.com/Luciani-Group. Immunofluorescent staining images were visualized using Imaris 10.1.0 (Oxford Instruments, Abingdon, UK). Confocal microscopy images are displayed as maximum intensity projections of the z-stack.

### 4.10. Statistical analysis

All results are reported as mean ± standard deviation (SD) of three independent replicates unless stated otherwise. Statistical analysis was performed in GraphPad Prism 10.3.0 for Windows (GraphPad Software, San Diego, USA). Pairs were compared using the Mann-Whitney test. For experiments with three or more groups, the Kruskal-Wallis test with Dunn’s multiple comparisons was used. To check the significance of differences between different formats, each group (monolayers, FN-silk networks, or spheroids) was compared to every other group. To check significance of differences between treatment groups and a control, each treatment group was compared to the control. Significance values were chosen as *p<0.1; **p<0.05.

### Acronyms

*α*-SMA *α* smooth muscle actin

12Z A human endometriotic epithelial cell line

2D Two-dimensional

3D Three-dimensional

A Alanine

a.u. arbitrary units

ANOVA Analysis of Variance

ATP Adenosine triphosphate

BSA Bovine serum albumin

C Cysteine

CCK-8 Cell Counting Kit-8

cDNA Complementary

DNA D Aspartic acid

D-PBS Dulbecco’s PBS

DAPI 4′,6-diamidino-2-phenylindole

DMEM Dulbecco’s Modified Eagle Medium

DMSO Dimethyl sulfoxide

ECM Extracellular matrix

EdU 5-ethynyl-2’-deoxyuridine

EpCAM Epithelial cell adhesion molecule

G Glycine

HCl Hydrochloric acid Ki67 Antigen Kiel 76

P Proline

p Passage number

PBS Phosphate-buffered saline

PFA Paraformaldehyde

PFD Pirfenidone R Arginine

R-Smads Receptor-regulated Smad proteins

RT-qPCR Quantitative reverse transcription polymerase chain reaction

S Serine

SB Sudan Black

SD Standard deviation

SDG Sustainable Development Goals

SDS Sodium dodecyl sulfate

Smad Suppressor of mothers against decapentaplegic

STR Highly polymorphic short tandem repeat locus

T Threonine

T-HESC A human endometrial stromal cell line

TGF-β Transforming growth factor β

V Valine

## Acknowledgements

We wish to thank Spiber Technologies AB for generously providing the FN-silk protein. ST wishes to thank Dr. Simone Aleandri for helpful discussions on statistical analysis and Astrid Källén, Savvina Gkouma, Dr. Kelly Blust, Dr. Inês Marquez, Dr. Yury Belyaev, Dr. Cristina Zivko, Dr. Thomas Tapmeier, Dr. Assad Riaz, Dr. Anna Stejskalová, and Dr. Shannon Hawkins for sharing their expertise on cell culture methods. ST wishes to thank Dr. Jiri Nohava from Anton Paar TriTec SA for the Bioindenter measurements. ST, MB and PL thank Prof. Dr. Oliver Mühlemann and Nicole Kleinschmidt for providing resources for RT-qPCR analysis. ST acknowledges a Short Travel Grant for (Post)Docs from the University of Bern and the Microscopy Imaging Center at the University of Bern. PL acknowledges the Swiss National Science Foundation, grant number 215227, for funding part of this project. Some figures were created with BioRender.com.

## CRediT authorship statement

ST – conceptualization (equal), data curation (equal), formal analysis (equal), funding acquisition (supporting), investigation (equal), methodology (lead), validation (supporting), visualization (lead), writing – original draft (lead), writing – review & editing (equal) MCB – data curation (equal), formal analysis (equal), investigation (equal), methodology (supporting), validation (lead), writing – review & editing (equal) MW – methodology (supporting), supervision (supporting), writing – review & editing (equal) LAG – methodology (supporting), supervision (supporting), writing – review & editing (equal) MH – funding acquisition (supporting), project administration (supporting), resources (supporting), supervision (supporting), writing – review & editing (equal) PL – conceptualization (equal), funding acquisition (lead), project administration (lead), resources (lead), supervision (lead), writing – review & editing (equal)

## Data availability statement

Data available on request from the authors.

