## Supplementary Information for "A Fibronectin (FN)-Silk 3D Cell Culture Model as a Screening Tool for Repurposed Antifibrotic Drug Candidates for Endometriosis"

**Characterization of T-HESC and 12Z monolayers**

**Figure S.1** shows monolayers of the T-HESC line (**Figure S.1A**) and 12Z cell lines (**Figure S.1B**), with actin filaments of the cytoskeleton shown in red and nuclei shown in blue. In monolayers, these cells reveal distinct morphologies consistent with their cell type. T- HESCs are large and clearly spindle-shaped with actin filaments of the cytoskeleton oriented along the longitudinal axis of the cell (**Figure S.1A**). 12Z cells are smaller and have lost their epithelial phenotype, showing a more spindle-like shape (**Figure S.1B, C**). This phenotype is consistent with 12Z cells' expression of N-cadherin reported in [S1], indicating that cells have experienced a degree of epithelial-to-mesenchymal transition.[S2] **Figure S.1C** shows that T-HESCs have a larger area at 7287 ± 3729 μm$^2$ cell$^{-1}$ compared to 12Z cells at 5734 ± 3123 μm$^2$ cell$^{-1}$ (mean ± SD, Supporting Information **Figure S.2**). 12Z cells have double the metabolic activity of T-HESCs (**Figure S.1D**), consistent with their shorter doubling time.

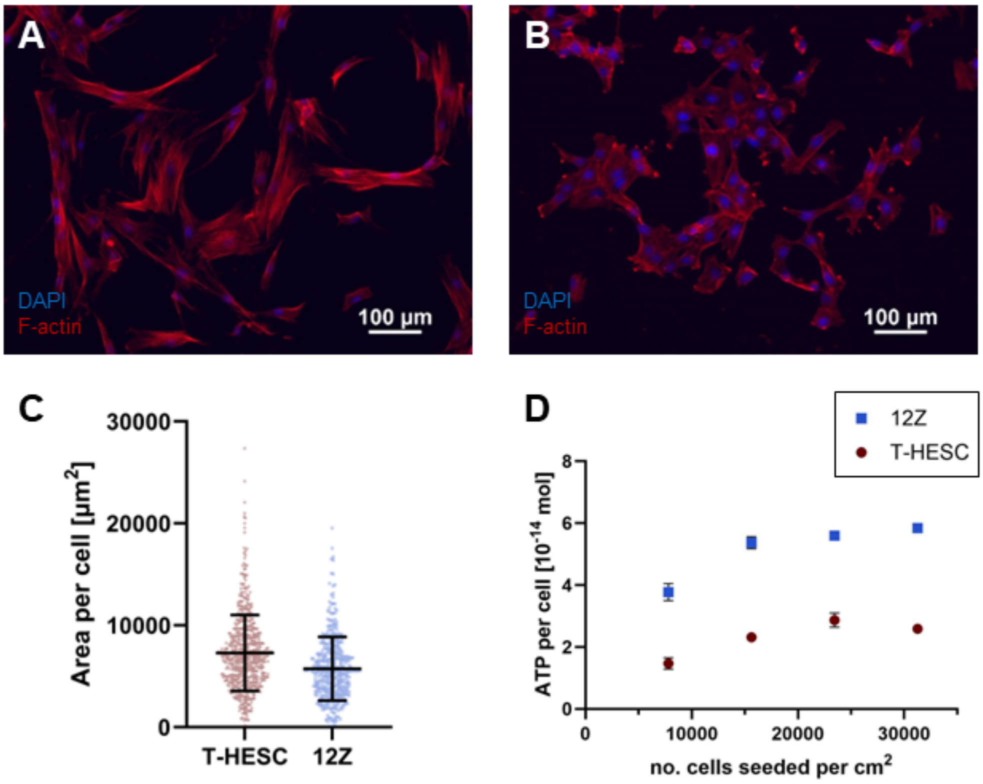


**Figure S.1.** Endometriotic cells cultured in monolayers. **A** T-HESC and **B** 12Z cell monolayers stained with phalloidin 594 (red, F-actin filaments of the cell cytoskeleton) and DAPI (blue, nuclei). **C** Area per cell of T-HESC and 12Z cells (mean ± SD from 589 T- HESCs and 399 12Z cells in a monolayer), measured using cell segmentation (Supporting Information **Figure S.2**). **D** ATP per cell against monolayer seeding density of T-HESCs and 12Z cells (mean ± SD from N=3 independent experiments, each with n=3 technical replicates).

**Measurement of area per cell**

Cell segmentation was performed using the CellPose algorithm[3] on **Figure S.2A** for T-HESCs and **Figure S.2B** for 12Z cells. Images were analyzed by predicting cell outlines (**Figure S.2C, D**), creating a mask (**Figure S.2E, F**), and computing the cell pose (**Figure S.2G, H**) as described in [3]. Code is provided on GitHub (https://github.com/Luciani-Group/).

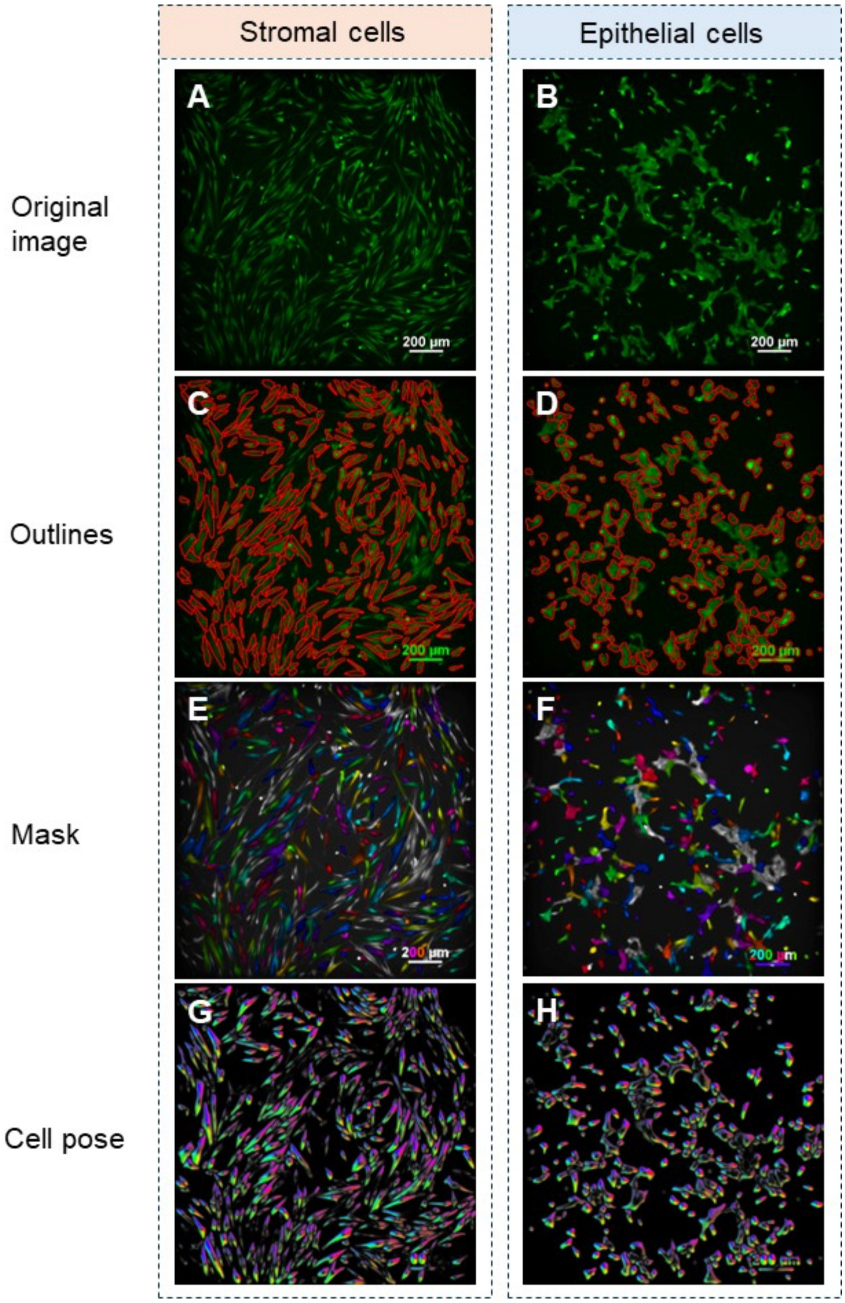


**Figure S.2.** Cell segmentation analysis of T-HESC and 12Z cell monolayers. **A** T-HESC and **B** 12Z cell monolayers with the cytoplasm stained with calcein AM. Predicted cell outlines of **C** T-HESC and **D** 12Z cells. Predicted mask of **E** T-HESC and **F** 12Z cells. Cell pose of **G** T-HESC and **H** 12Z cells.

### Detachment of cells from FN-silk networks

Attempts were made to detach T-HESC and 12Z cells from FN-silk networks at day 7 of the culture period when cells were expected to have the strongest adhesion to the FN-silk matrix. FN-silk networks were washed with PBS three times, incubated in each respective detachment medium in a humidified incubator at 37 °C, 5% $CO_2$, and disrupted mechanically. Cells in the supernatant were stained using Trypan blue and counted using a Neubauer plate. **Figure S.3A and S.3B** show that neither T-HESC nor 12Z cells could be recovered at high levels from the FN-silk networks, even under relatively harsh conditions of 20 min in TrypLE enzyme. **Figure S.3C** shows nuclei of T-HESCs that remained stuck to the FN-silk matrix after incubation of the network in TrypLE for 20 min. These results suggest that T-HESCs and, to a lesser extent, 12Z cells form strong cell-matrix interactions with the FN-silk.

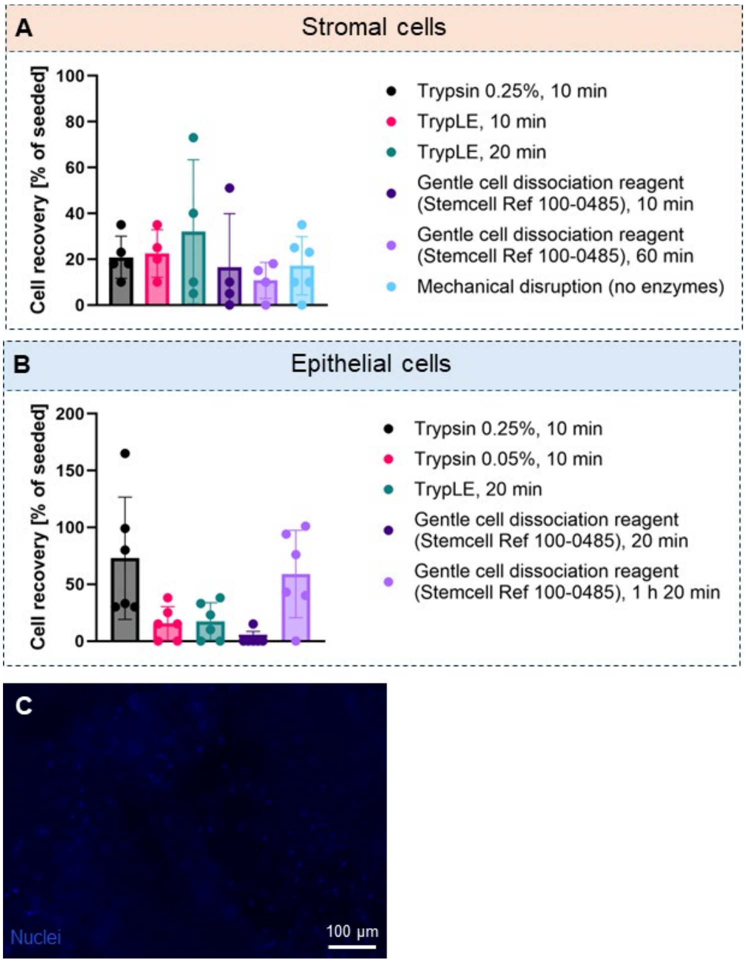


**Figure S.3.** Detachment of **A** T-HESC and **B** 12Z cells from FN-silk networks at culture day 7 using various enzymatic and nonenzymatic methods. Fraction of cells recovered relative to number of cells seeded (individual data points and mean ± SD from N=1 independent experiment with n=4−6 networks per condition (i.e., technical replicates). **C** Nuclei remaining on a T-HESC FN-silk network after incubation with TrypLE for 20 min.

### Cell proliferation in 3D: comparison of CellTiter Glo assay to manual count

To validate the CellTiter Glo® Cell 3D Viability Assay on FN-silk networks, and to determine how well ATP levels correlate with cell number and hence cell proliferation, the luminescence intensity measured using the CellTiter Glo 3D Cell Viability Assay was compared to the number of cells in the 3D system, measured by manual counting using a Neubauer plate.

3D systems (FN-silk networks and spheroids) were seeded as described in Section 2.2.2, except that stromal cell 3D systems were seeded at 20,000 cells per unit. The cell number/metabolic activity was measured at culture days 1, 4 and 7. At each time point, 10-11 3D units (FN-silk networks or spheroids) were used for manual counting and 4-5 3D units were used for the CellTiter Glo 3D Viability Assay. For manual counting, cells were detached from the FN-silk matrix as shown in **Figure S.4A**. To detach cells from the FN-silk matrix, FN-silk networks were washed in Dulbecco's phosphate buffered saline (D-PBS) containing calcium and magnesium (Gibco™ Ref 14040117, Grand Island, NY, USA) three times and incubated in Trypsin-EDTA 0.5% (Gibco™ 15400054, Canada) for 30 min in a humidified incubator at 37 °C, 5% $CO_2$. Complete growth medium was added to inactivate the Trypsin enzyme and 3D systems were pipetted up and down 20 times to allow shear forces to mechanically disrupt the 3D structure. Cells were stained with a 0.4% Trypan blue solution (Merck Ref T8154, Switzerland) and counted manually using a Neubauer plate, which revealed that cells had completely detached from fragments of FN-silk (images not shown). The expansion index was calculated using Equation 2.

$$Expansion\ index\ on\ day\ d = \frac{Number\ of\ cells\ on\ day\ d}{Number\ of\ cells\ on\ day\ 1} \quad (2)$$

The CellTiter Glo 3D Cell Viability Assay was performed as described in the Materials & Methods. The luminescence was normalized to the luminescence on day 1. **Figure S.4B** shows the expansion index of epithelial cells cultured in FN-silk networks and spheroids, assessed by a manual count. A comparison to **Figure S.4D**, which shows the same cell systems measured using luminescence, shows a similar trend, albeit with different magnitudes. For FN-silk networks, cells proliferated from culture day 1 to day 4, with the cell number increasing by a factor of 2.4 ± 0.5 compared to the cell number on day 1 (**Figure S.4B**). The corresponding luminescence had only increased by a factor of 1.5 ± 0.3 at day 4 relative to day 1 (**Figure S.4D**). By day 7, the number of cells was 2.2 ± 0.5 relative to day 1 (**Figure S.4B**), while the relative luminescence was 1.1 ± 0.4 (**Figure S.4D**).

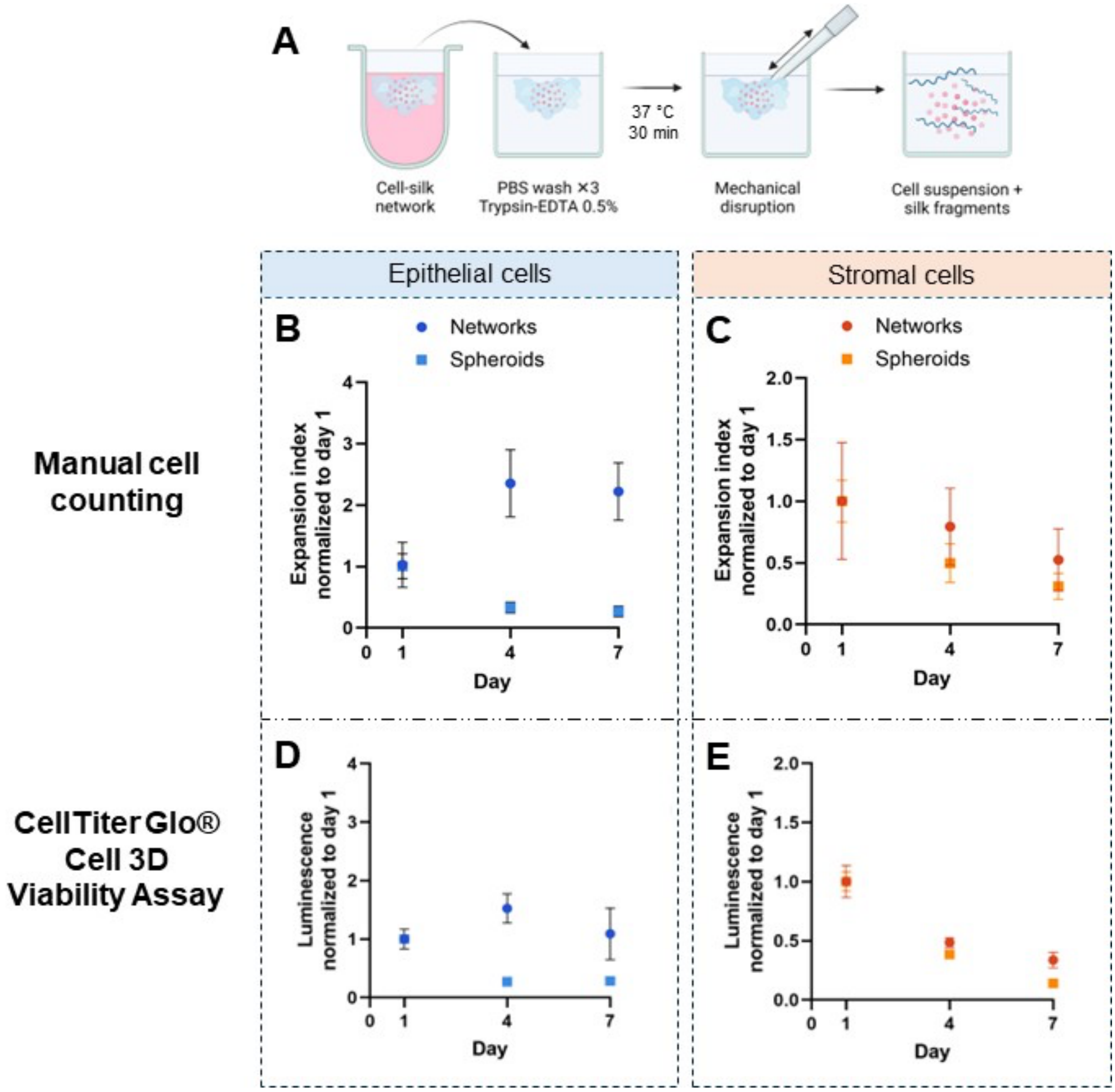


**Figure S.4.** Validation of CellTiter Glo Cell 3D Viability Assay by comparison to manual counting of cells. **A** Method to detach cells from FN-silk network to allow manual counting. Expansion index of **B** epithelial 12Z and **C** stromal T-HESCs cultured in FN-silk networks and spheroids evaluated by manual cell counting. Relative luminescence of **D** epithelial 12Z and **E** stromal T-HESCs cultured in FN-silk networks and spheroids evaluated using the CellTiter Glo Cell 3D Viability Assay. Mean ± SD from N=1 independent experiment with n=10 or n=11 technical replicates for manual counting and n=4 or n=5 technical replicates for the CellTiter Glo Cell 3D Viability Assay.

A possible explanation for the discrepancy between the cell number and metabolic activity is that cells were proliferating, but the metabolic activity per cell decreased. Alternatively, cells may have been in a cycle phase that is associated with lower metabolic activity, although this contradicts recent evidence from other cell types suggesting that mitosis is associated with increased mitochondrial metabolism.[4] For epithelial cell spheroids, the expansion index was at

0.3 ± 0.1 at days 4 and 7 (**Figure S.4B**), while the relative luminescence was lower than day 1 by a factor of 0.28 ± 0.02 at days 4 and 7 (**Figure S.4D**), revealing similar growth trends using both methods. For stromal cells, both FN-silk networks and spheroids showed a reduction in cell proliferation/metabolic activity because they were seeded at 20,000 cells per unit, which was later found to be too high to allow stromal cells to grow in FN-silk networks. Comparing **Figures S.4C and E** shows that manual counting and metabolic activity give very similar results in terms of cell growth. Overall, manual counting showed higher variability than the luminescence-based CellTiter Glo Cell 3D Viability assay, which may be because a larger number of technical replicates was included, or because manual counting was performed at a cell concentration of $0.5-1.0\times10^5$ cells $mL^{-1}$, which is below the lower limit of the accurate range of the Neubauer plate method of $2.5\times10^5$ cells $mL^{-1}$. Overall, the cell counting and luminescence assay give similar results for three of four systems (epithelial spheroids, stromal FN-silk networks and stromal spheroids). For the fourth system (12Z FN-silk networks), the luminescence assay tended to underestimate cell proliferation compared to the manual counting method.

**Stromal cell FN-silk networks seeded at lower cell densities**

Stromal cells were seeded on FN-silk networks at lower seeding densities to assess whether the reduction in metabolic activity observed post-day 4 was attributable to the FN-silk matrix being full of cells. Stromal cell FN-silk networks were seeded as described in the Materials & Methods, except that cells were seeded at densities of 5000 and 2500 cells per network. Metabolic activity was evaluated using the CellTiter Glo Cell 3D Viability Assay as described in the Materials & Methods. **Figure S.5** shows that all seeding densities have in increase in cell metabolic activity from day 1 to day 4, followed up a decrease in metabolic activity from day 4 to day 7. This suggests that the reduction in metabolic activity observed post-day 4 is not attributable to the silk matrix being full of cells.

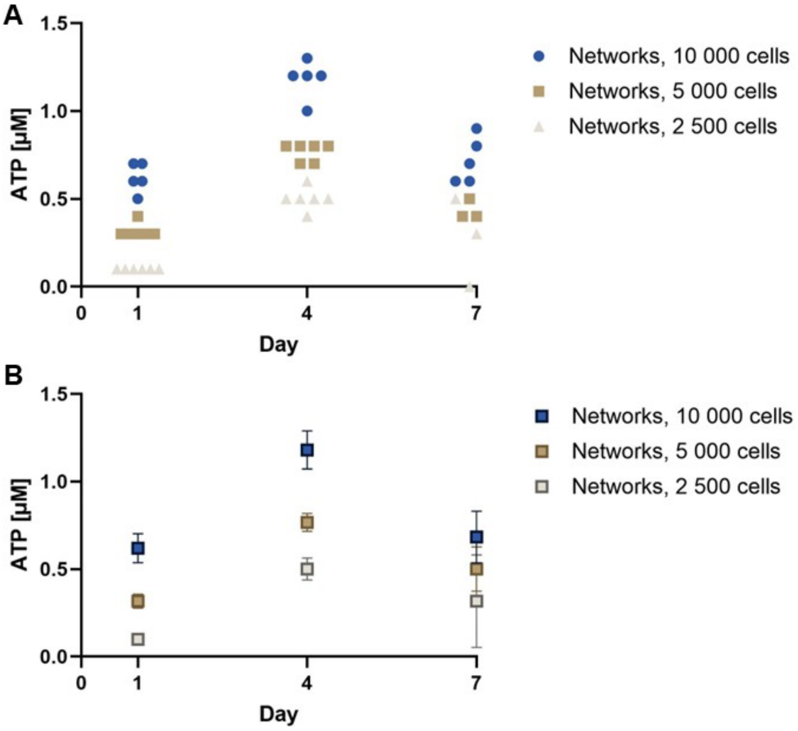


**Figure S.5.** Metabolic activity of stromal T-HESC FN-silk networks at lower seeding densities. **A** Individual data points and **B** mean ± SD from N=1 independent experiment with n = 3−6 FN-silk networks per time point (i.e., technical replicates).

**Morphology of FN-silk networks**

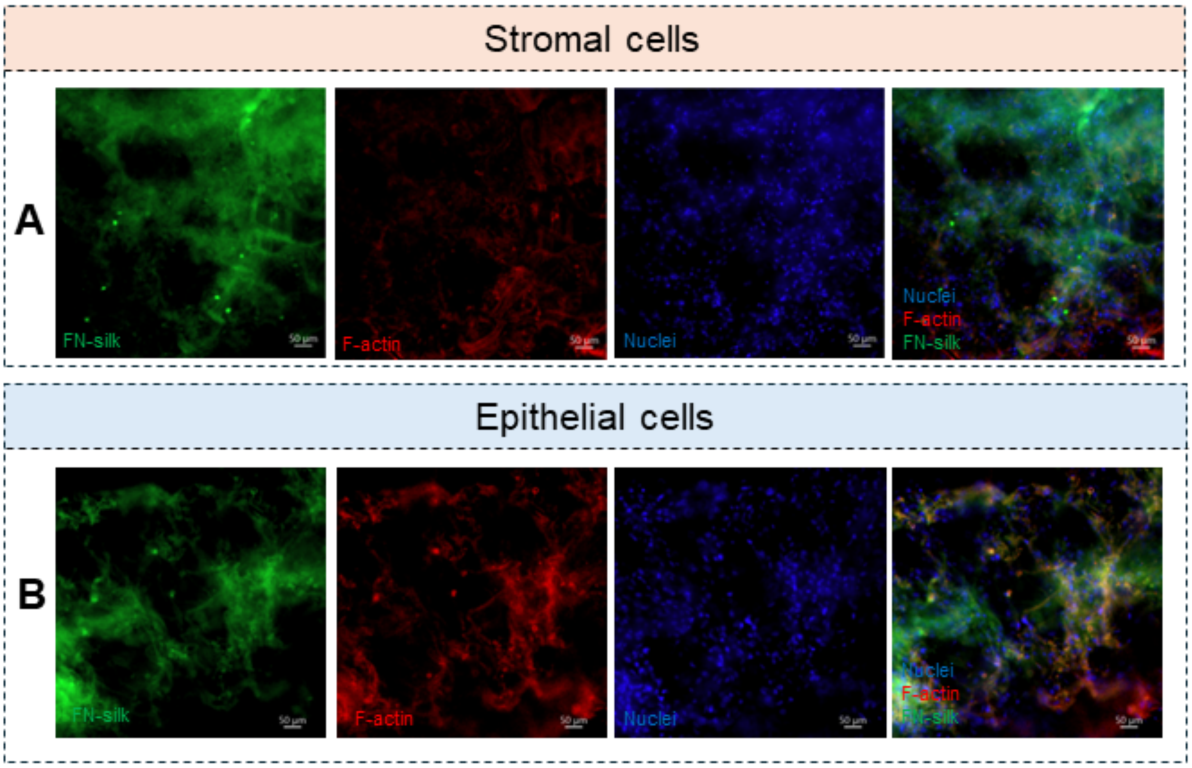


**Figure S.6.** Close up of **A** T-HESC FN-silk networks at culture day 7. **B** Close up of 12Z cell FN-silk networks on culture day 1. Green channel: FN-silk. Scale bars indicate 50 µm.

### Effect of pirfenidone on metabolic activity of T-HESC and 12Z monolayers

We assessed the effect of pirfenidone concentration on the viability of stromal T-HESCs and epithelial 12Z cells in monolayers using a CCK-8 assay, as described in the Materials & Methods. For both cell types, viability remained close to untreated cells at pirfenidone concentrations of up to 1 mg $mL^{-1}$, while pirfenidone concentrations above 1 mg $mL^{-1}$ reduced viability (**Figure S.7**).

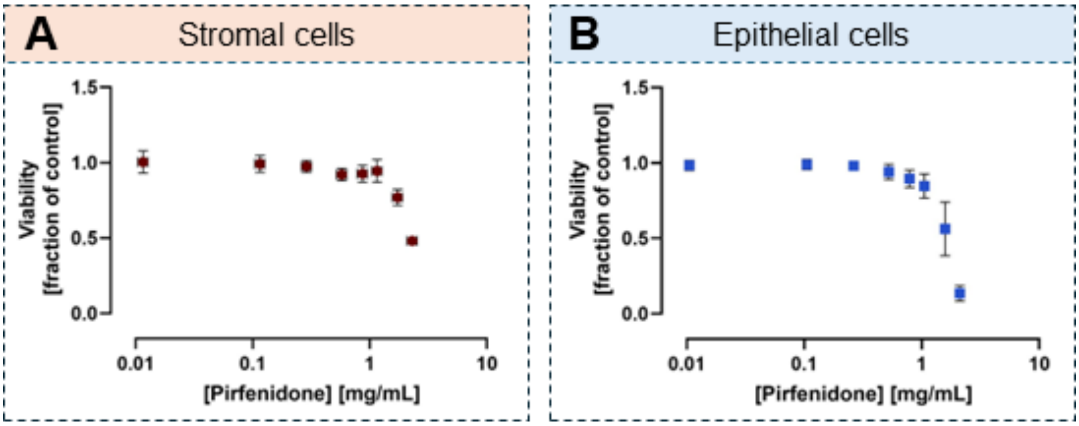


**Figure S.7.** Effect of pirfenidone concentration on the viability of **A** stromal T-HESC and **B** epithelial 12Z cell monolayers treated for 24 h (mean ± SD from N=3 independent experiments with n=3−6 technical replicates each).

### Cell line authentication

**Table S.1** shows a summary table of the STR profiling on the T-HESC sample. A 100% match between the submitted sample and database alleles was found. **Figure S.8** shows the electropherogram the T-HESC sample. **Table S.2** shows a summary table of the STR profiling on the 12Z sample. A 92.5% match between the submitted sample and database alleles was found. **Figure S.9** shows the electropherogram of the 12Z sample. Differences were found in D8S1179 and FGA loci. For the D8S1179 locus, the database shows 10 while the tested sample showed 9/10. According to abm, the supplier of the cell line, allele 9 is a typical stutter peak and this difference is negligible. For the FGA locus, the submitted sample showed 22/23 but the database shows 23/24. According to abm, this may be from mutations that accumulated during passaging the cells or from test variation. With this information, the submitted sample showed a 96% match with the 12Z cell line, the only difference being in the FGA locus.

**Table S.1.** Summary of stromal T-HESC sample and database STR profiles

| Locus | Chromosomal location | Core STR marker | Sample alleles | Database alleles |
|---|---|---|---|---|
| D3S1358 | Chr03 | | 16/17 | N/A |
| TH01 | Chr11 | Yes | 7/8 | 7/8 |
| D21S11 | Chr21 | | 30/31.2 | N/A |
| D18S51 | Chr18 | | 15/16 | N/A |
| Penta_E | Chr15 | | 5/7 | N/A |
| D5S818 | Chr05 | Yes | 8 | 8 |
| D13S317 | Chr13 | Yes | 12/13 | 12/13 |
| D7S820 | Chr07 | Yes | 10 | 10 |
| D16S539 | Chr16 | Yes | 12/14 | 12/14 |
| CSF1PO | Chr05 | Yes | 10/12 | 10/12 |
| Penta_D | Chr21 | | 10/11 | N/A |
| AMEL | X/Y | Yes | X | X |
| vWA | Chr12 | Yes | 16/18 | 16/18 |
| D8S1179 | Chr08 | | 14/16 | N/A |
| TPOX | Chr2 | Yes | 9/11 | 9/11 |
| FGA | Chr04 | | 20/21 | N/A |

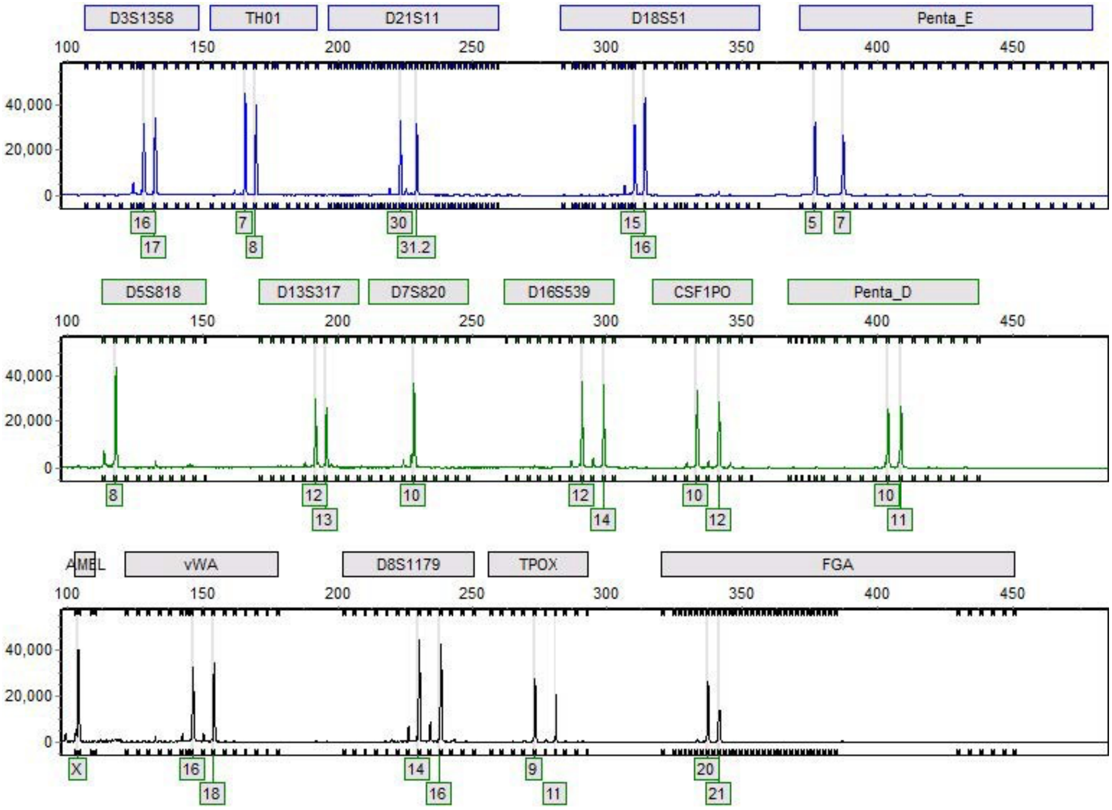


**Figure S.8.** Electropherogram of T-HESC sample

**Table S.2.** Summary of epithelial 12Z sample and database STR profiles

| Locus | Chromosomal location | Core STR marker | Sample alleles | Database alleles |
|---|---|---|---|---|
| D3S1358 | Chr03 | | 14/17 | 14/17 |
| TH01 | Chr11 | Yes | 9/9.3 | 9/9.3 |
| D21S11 | Chr21 | | 28/30.2 | 28/30.2 |
| D18S51 | Chr18 | | 13/15 | 13/15 |
| Penta_E | Chr15 | | 11/12 | N/A |
| D5S818 | Chr05 | Yes | 12 | 12 |
| D13S317 | Chr13 | Yes | 12/13 | 12/13 |
| D7S820 | Chr07 | Yes | 9/12 | 9/12 |
| D16S539 | Chr16 | Yes | 13 | 13 |
| CSF1PO | Chr05 | Yes | 11/13 | 11/13 |
| Penta_D | Chr21 | | 9/12 | N/A |
| AMEL | X/Y | Yes | X | X |
| vWA | Chr12 | Yes | 17 | 17 |
| D8S1179 | Chr08 | | 9/10 | 10 |
| TPOX | Chr2 | Yes | 11/12 | 11/12 |
| FGA | Chr04 | | 22/23 | 23/24 |

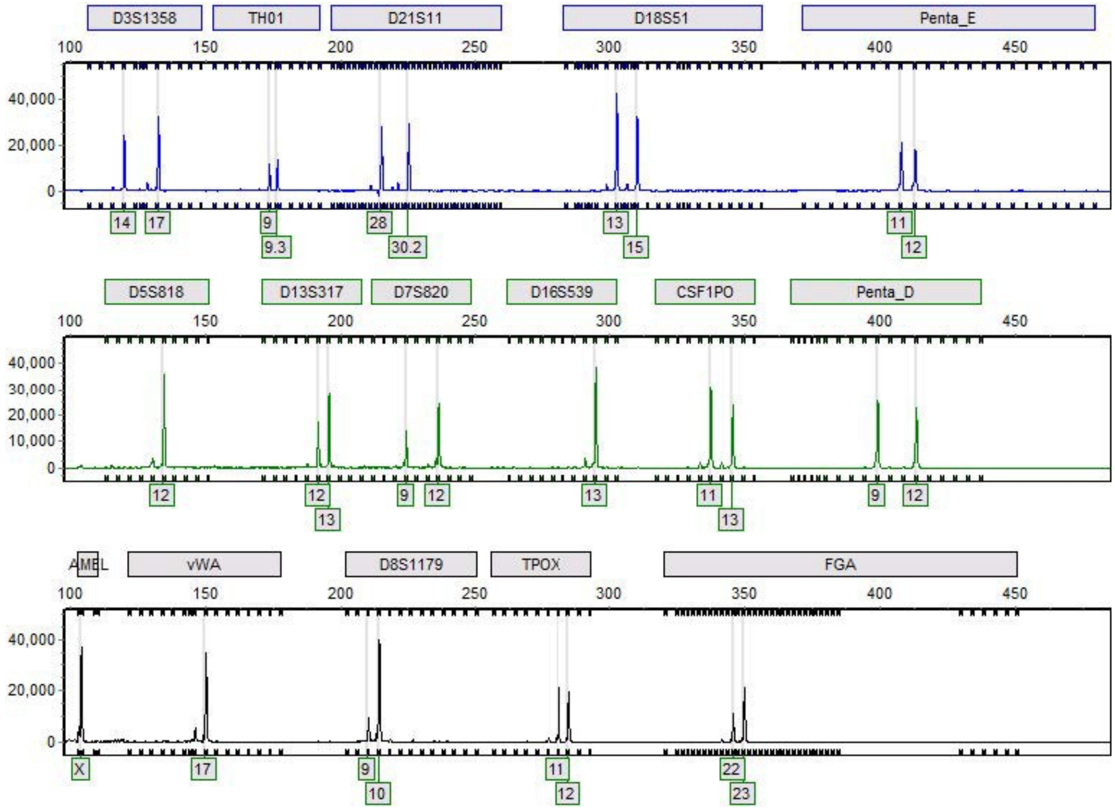


**Figure S.9.** Electropherogram of 12Z cell sample

**Immunofluorescent staining control**

For immunofluorescent staining, a control sample was stained with secondary antibody only (no primary antibody) to check for unspecific binding of the secondary antibody. **Figure S.11** shows low fluorescence in the green channel, which confirms that green channel fluorescence observed in samples stained with primary antibody is attributable to expression of the respective proteins, rather than unspecific binding of the secondary antibody.

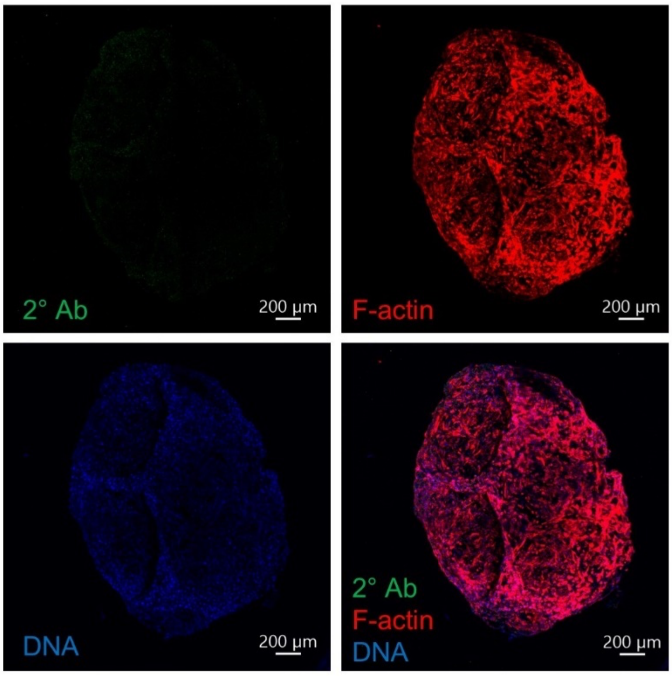


**Figure S.11:** Control immunofluorescent staining: T-HESC FN-silk network stained with secondary antibody, but no primary antibody. F-actin (phalloidin 594) and nuclei (DAPI) counterstained. Single channel images and overlay. Scale bars indicate 200 µm.

### RT-qPCR primers

Primer sequences and amplification efficiencies are provided in **Table S.3.**

**Table S.3.** Primer pair sequences used for RT-qPCR: forward and reverse sequences, amplification efficiency *E*, slope *n* of the linear regression of $\Delta C_t(gene\text{-}GAPDH)$ vs. log(cDNA concentration), and source.

| Gene | Sequence (5'-3') | *E* | *n* | Source |
|---|---|---|---|---|
| *GAPDH* fwd | GAG TCA ACG GAT TTG GTC G | 86% | | [9] |
| *GAPDH* rev | GAG GTC AAT GAA GGG GTC AT | | | |
| *COL1A1* fwd | GTT CAG TTT GGG TTG CTT GTC T | 80% | –1.0180 | Thermo Fisher Scientific Hs00521314_CE |
| *COL1A1* rev | CCT GCC CAT CGA TGT G | | | |
| *ACTA2* fwd | CTA TGA GGG CTA TGC CTT GCC | 89% | –0.3700 | [6] |
| *ACTA2* rev | GCT CAG CAG TAG TAA CGA AGG | | | |
| *Fn-1* fwd | AGG AAG CCG AGG TTT TAA CTG | 81% | –0.1807 | [6] |
| *Fn-1* rev | AGG ACG CTC ATA AGT GTC ACC | | | |
| *Smad3* fwd | TGG ACG CAG GTT CTC CAA AC | 82% | –0.1082 | [6] |
| *Smad3* rev | CCG GCT CGC AGT AGG TAA C | | | |
| *CDH1* fwd | GCC TCC TGA AAA GAG AGT GGA AG | 89% | –0.1799 | OriGene NM004360 |
| *CDH1* rev | TGG CAG TGT CTC TCC AAA TCC G | | | |
| *CDH2* fwd | CCT CCA GAG TTT ACT GCC ATG AC | 82% | –0.1159 | OriGene NM001792 |
| *CDH2* rev | GTA GGA TCT CCG CCA CTG ATT C | | | |
| *SNAI1* fwd | CAC TAT GCC GCG CTC TTT C | 82% | –0.1776 | [10] |
| *SNAI1* rev | GGT CGT AGG GCT GCT GGA A | | | |
| *SNAI2* fwd | ATC TGC GGC AAG GCG TTT TCC A | 82% | –0.1008 | OriGene HP206658 |
| *SNAI2* rev | GAG CCC TCA GAT TTG ACC TGT C | | | |

### Suitability of GAPDH as a reference gene

GAPDH was found to be a suitable housekeeping gene, as its expression was unaffected by TGF-β1 treatment at a concentration of 10 ng $mL^{-1}$ for 24 h compared to vehicle treatment. GAPDH $C_t$ values of TGF-β1 and vehicle-treated T-HESC and 12Z cells are shown in **Figure S.12A and S.12B**, respectively.

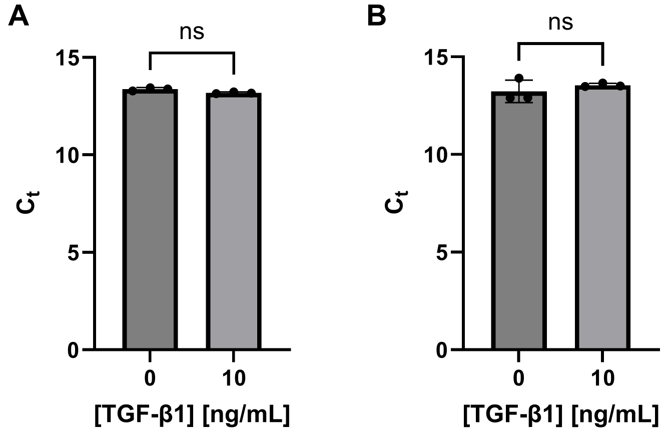


**Figure S.12.** Effect of TGF-β1 treatment on the $C_t$ value of the housekeeping gene GAPDH for **A** stromal T-HESC and **B** epithelial 12Z cells (mean ± SD from n=3 technical replicates).

### References to the Supporting Information

[S1] A. Zeitvogel, R. Baumann, A. Starzinski-Powitz, *Am. J. Pathol.* **2001**, *159*, 1839.

[S2] L. Ma, T. Andrieu, B. McKinnon, L. Duempelmann, R.-W. Peng, C. Wotzkow, C. Müller, M. D. Mueller, *J. Steroid Biochem. Mol. Biol.* **2021**, *212*, 105943.

[S3] C. Stringer, T. Wang, M. Michaelos, M. Pachitariu, *Nat. Methods* **2021**, *18*, 100.

[S4] F. F. Diehl, K. M. Sapp, M. G. Vander Heiden, *Trends Cell Biol.* **2024**, *34*, 136.

[S5] M. G. Ibrahim, M. Sillem, J. Plendl, E. T. Taube, A. Schüring, M. Götte, V. Chiantera, J. Sehouli, S. Mechsner, *Arch. Gynecol. Obstet.* **2019**, *299*, 489.

[S6] Y. Dai, L. Xin, S. Hu, S. Xu, D. Huang, X. Jin, J. Chen, R. W. S. Chan, E. H. Y. Ng, W. S. B. Yeung, L. Ma, S. Zhang, *Regen. Biomater.* **2023**, *10*, rbad080.

[S7] M. Song, L. Ma, Y. Zhu, H. Gao, R. Hu, *Sci. Rep.* **2024**, *14*, 8321.

[S8] S. Matsuzaki, J.-L. Pouly, M. Canis, *Sci. Rep.* **2020**, *10*, 9467.

[S9] A. A. Ganguin, I. Skorup, S. Streb, A. Othman, P. Luciani, *Adv. Healthc. Mater.* **2023**, *12*, 2300811.

[S10] H. Wang, E. Chirshev, N. Hojo, T. Suzuki, A. Bertucci, M. Pierce, C. Perry, R. Wang, J. Zink, C. A. Glackin, Y. J. Ioffe, J. J. Unternaehrer, *Cancers* **2021**, *13*, 1469.
